# Exploration of a European-centered strawberry diversity panel provides markers and candidate genes for the control of fruit quality traits

**DOI:** 10.1101/2024.03.04.583203

**Authors:** Alexandre Prohaska, Pol Rey-Serra, Johann Petit, Aurélie Petit, Justine Perrotte, Christophe Rothan, Béatrice Denoyes

## Abstract

Fruit quality traits are major breeding targets in cultivated strawberry (*Fragaria × ananassa*). Taking into account the requirements of both growers and consumers when selecting high quality cultivars is a real challenge. Here, we used an original diversity panel enriched with unique European accessions and the 50K FanaSNP array to highlight the evolution of strawberry diversity over the past 160 years, investigate the molecular basis of 12 major fruit quality traits by GWAS, and provide genetic markers for breeding. Results show that considerable improvements of key breeding targets including fruit weight, firmness, composition and appearance occurred simultaneously in European and American populations. Despite the high genetic diversity of our panel, we observed a drop in nucleotide diversity in certain chromosome regions, revealing the impact of selection. GWAS identified 71 associations with 11 quality traits and, while validating known associations (firmness, sugar), highlighted the predominance of new QTL, demonstrating the value of using untapped genetic resources. Three of the six selective sweeps detected are related to glossiness or skin resistance, two little-studied traits important for fruit attractiveness and, potentially, postharvest shelf-life. Moreover, major QTL for firmness, glossiness, skin resistance and susceptibility to bruising are found within a low diversity region of chromosome 3D. Stringent search for candidate genes underlying QTL uncovered strong candidates for fruit color, firmness, sugar and acid composition, glossiness and skin resistance. Overall, our study provides a potential avenue for extending shelf-life without compromising flavor and color as well as the genetic markers needed to achieve this goal.

## Introduction

Cultivated strawberry *(Fragaria* × *ananassa)*, the most widely consumed small fruit worldwide, results from spontaneous hybridization in botanical gardens in France in the 18^th^ century between two octoploid (2n = 8x = 56) species (*F. chiloensis* and *F. virginiana*) imported from the New World^1^. Since then, cultivated strawberry has been continuously improved through the introgression of alleles from wild progenitors creating an admixed population of interspecific hybrid lineages^2,3,4^. Recurrent hybridization contributed to maintain genetic diversity in the domesticated populations^4^. However, lower genetic diversity and heterozygosity can be observed in highly structured populations, which nevertheless show considerably improved yield, fruit weight and firmness^5^. Current efforts, triggered by consumer demand for sweet and highly-flavored strawberries^6,7^, are aimed at improving the sensory and nutritional quality of the fruit, for example color^8^ and flavor^9^. Another area for improvement is the extension of the storage period and the reduction of post-harvest rot, both of which are linked to fruit firmness^7^ and little-explored fruit surface properties^10^. Several fruit quality traits can be easily manipulated using advanced technology, such as genome editing which has successfully been applied to create new alleles modifying various traits including fruit color, sweetness and aroma^7,11^. Other complex (e.g. fruit size) and/or little-studied (e.g. fruit glossiness) traits first require elucidation of their genetic architecture. Until recently, following initial studies^12,13^, the dissection of the genetic control of complex fruit quality traits in *F.*× *ananassa* has mainly been achieved by mapping Quantitative Trait Locus (QTL) on genetic linkage maps of bi-parental^14,15,16^ or multi-parental^17^ populations. Causal genetic variants have been identified for several QTL, leading to the design of genetic markers for marker-assisted selection (MAS) of strawberry varieties with, for example, improved fruit color^8,18^ and better aroma^9^.

Complexity of the allo-octoploid genome of *F.*x *ananassa*, where up to eight homeo-allelic forms of the same gene can be found^19^, has until recently hampered the mapping of QTLs on a given chromosome. Whole genome sequencing of *F.*x *ananassa*^1^ and, more recently, its progenitors^20^, showed that the four subgenomes of *F.* x *ananassa* are derived from the diploids *F. vesca* and *F. iinumae* and from two extinct species related to *F. iinumae*^20^. Genome sequence further enabled the design of a single nucleotide polymorphism (SNP) 50K array with selected chromosome-specific SNPs^21^ allowing the high-resolution mapping of QTLs. A recent breakthrough has been the completion of haplotype-resolved genomes for five genotypes of *F.*× *ananassa*^9,22,23,24,25^. These developments make it possible to exploit strawberry diversity through genome wide association studies (GWAS), which scans the genome for significant associations between genetic markers and the trait studied^2^. It thus can help unveil beneficial alleles through the exploration and characterization of large strawberry genetic resources, which display a wide genetic and phenotypic diversity^4,26,27,28^. So far, GWAS has been done on collections mostly centered on North America strawberry populations^4,28^, which enabled the discovery of major QTLs controlling fruit weight, firmness, sweetness and aroma^4,9,28,29,30^. It would certainly benefit from the exploration of other less well-characterized but original populations found in Europe^31^, which is one of the most active strawberry breeding centers^6^.

In this study, we explored by GWAS the genetic architecture of fruit quality in *F.*× *ananassa*. To this end, we used the unexploited genetic diversity found in traditional and modern European varieties, in comparison with the better described diversity of North American varieties and some Asian genotypes. Our results are consistent with recent insights into the evolution of modern strawberry varieties and detected major QTL recently described, for fruit firmness for example. Moreover, we detected new QTL for most of the 12 fruit quality traits studied and the underlying candidate genes (CG). An example of this are the QTL and strong CG for the little-explored glossy trait, which underpins the shiny appearance of all modern strawberry varieties and was found to co-localize with a skin resistance trait. Our results therefore highlight the richness of European collections as a source of genetic diversity for strawberry breeding.

## Results

### Population structure and genetic diversity of the diversity panel of cultivated strawberry

We analyzed a germplasm diversity panel comprising 223 accessions of cultivated strawberries (*F.*× *ananassa*) available at Invenio (South-West France) (Table S1). Unlike the main diversity panels studied to date, where the bulk of the panel was constituted by North American accessions^4,27^, our panel was mostly composed of European accessions from several countries, with French cultivars being by far the most represented (96 accessions). In addition, the panel comprised representative cultivars from North America including California and Florida, Japan and other breeding programs around the world (Fig. 1A, Table S1). Many cultivars were released between 1990s and nowadays, but the panel also accurately covered the whole modern breeding period (>1950s) and the early stages of strawberry breeding, with cultivars reaching as far as the beginning of the 19th century (Fig. S1). Thirty-two accessions from this panel were common with those from the study by Horvath et al. (2011)^31^. Accessions from the diversity panel were genotyped with the 50K FanaSNP array^21^. A total of 38,120 SNPs was retained after filtering for minor allelic frequency (MAF) (< 5%) and missing data (> 3%).

**Figure 1.**
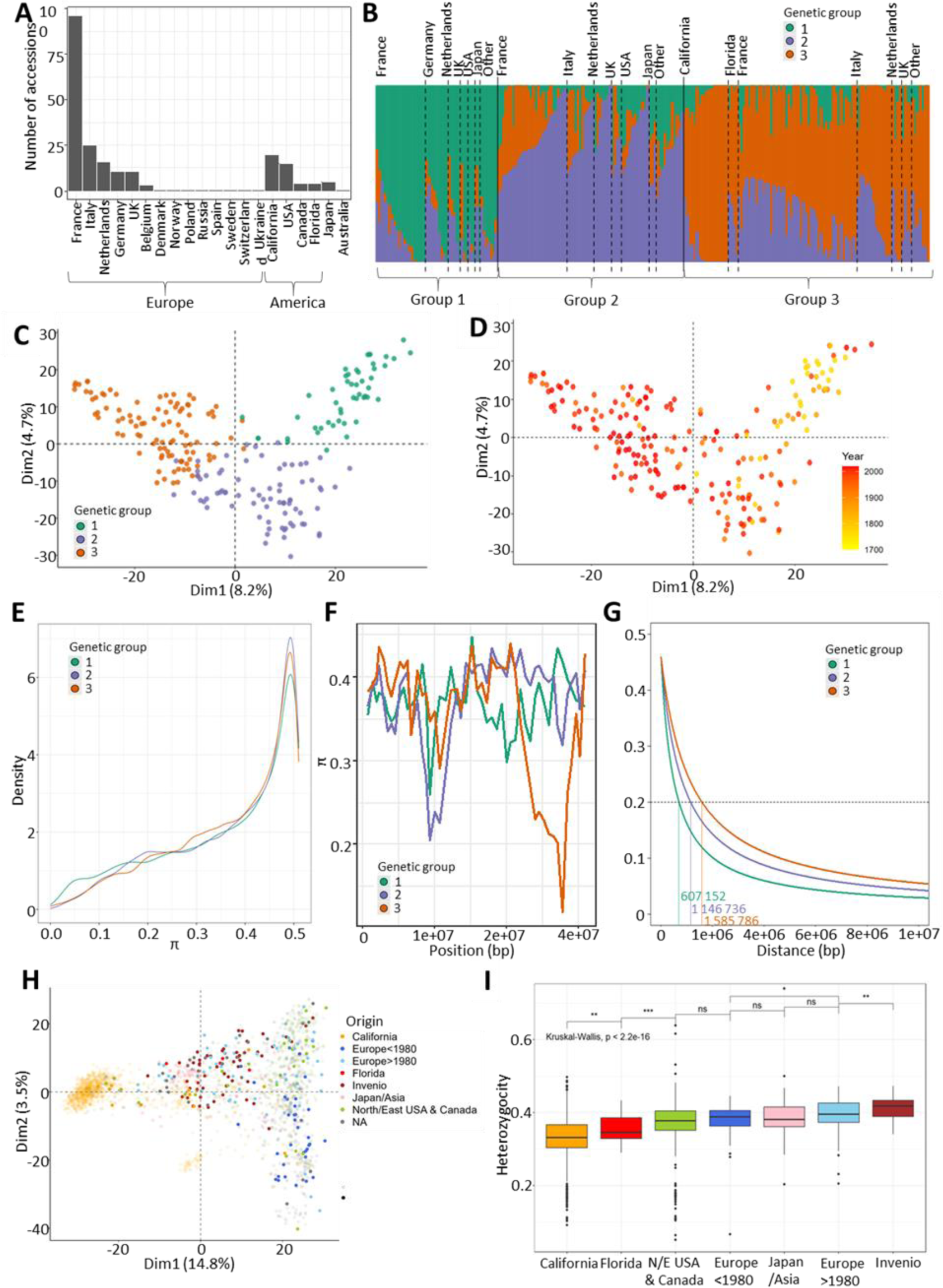
Genetic diversity of the panel. (A) Distribution of the geographical origin of the 223 accessions. (B) Structure barplot representing each genotype (bars) by its percentage of affiliation to each of the three genetic groups according to the STRUCTURE analysis. Individuals are sorted by genetic groups and geographical origins. (C,D) Principal Component Analysis of 38 120 SNP markers. Each accession (dot) is colored by its genetic group (C) or year of release (D). (E) Nucleotide diversity (π) distributions in windows of 400 kb across each genetic group. (F) π chromosome-wide estimates for each genetic group for 400kb windows across the chromosome 3D. (G) Linkage disequilibrium (LD) decay along chromosome 1A. The dashed line represents the LD decay at r^2^=0.2. (H) Distribution of the Invenio panel (filled dots) among 1 569 genotypes (shaded dots) studied in Hardigan et al. (2021B)^4^ and 539 genotypes studied in Zurn et al. (2022)^27^ (shaded dots) with 3215 SNP markers. Accessions are colored according to geographical/breeding origin. (I) Heterozygosity coefficients across different geographical/breeding origins when combining accessions from the diversity panel and 1569 genotypes from Hardigan et al. (2021b)^4^. Genetic groups 1, 2 and 3 are colored in green, purple and orange, respectively. Groups 1, 2, 3: Heirloom & related, European mixed group and American & European mixed groups, respectively.

To explore relationships among the 223 accessions, we first evaluated the population structure with STRUCTURE software. We identified three distinct genetic clusters (Fig. 1B, Table S1). Group 1 (G1) includes most of the older European cultivars and their more recent relatives, as well as some old American cultivars (Fig. 1B). This group is hereafter named the Heirloom & related group. Group 2 (G2) clusters essentially European, as well as 14 American cultivars mostly from North-East America (Maryland, New-York and Canada) and three out of five Japanese cultivars (Fig. 1B) and was therefore named the European mixed group. Californian and Floridian cultivars, together with other European ones, have been identified in group 3 (G3) hereafter named the American & European mixed group. A large amount of admixture (< 70%) was observed for each group, with 108 out of 223 accessions split across more than one group, with most of the admixture spread between G2 and G3 (Fig. 1B).

To further investigate population structure, we performed principal component analysis (PCA) of the 223 accessions using the 38,120 SNP markers (Fig. 1C). The first two dimensions (PC1, PC2) explained 8.2% and 4.7% of the structural variance, respectively. The three genetic groups were positioned at each vertex of the crescent shape. PC1 also reflected the temporal separation between G1 and the other two groups when cultivars were displayed according to year of release (Fig. 1D). The phylogenetic analysis (Fig. S2) was consistent with the structure (Fig. 1B) and PCA (Figs 1C, 1D) analyses.

Genome-wide comparisons of nucleotide diversity (π) between genetic groups revealed no clear loss of genetic diversity from G1 to G2 and G3 (Fig. 1E). At the chromosome level, the distribution of nucleotide diversity among groups was uneven, with several genomic regions associated with significant enrichment or loss. Of notice, some regions were associated with a sharp reduction in diversity in G2 and/or G3 compared with G1, for example on chromosome 3D, from 23,233 kb to 29,635 kb (Fig. 1F, Fig. S3). Progression towards the most recent American and European cultivars also translated in local augmentation in linkage disequilibrium, where LD (at r² = 0.20) increased from an average distance of 802,184 bp for G1 to 1,073,213 bp for G2 and 1,253,777 bp for G3 (Fig. 1G, Fig. S4).

We then combined the SNP data from our diversity panel with those from ^4^ (University of California Davis, UCD panel) and ^27^ (United States Department of Agriculture, USDA panel) to position our collection in relation with these studies. The PCA of the combined data revealed that the Invenio collection largely overlapped the two USA collections, with the exception of the extreme end of the PC1 corresponding to the UCD program and a small group of genotypes representing probable introgressions of wild accessions into the Californian panel (Fig. 1H, Sup. Fig. 6). The University of Florida (FL) program was less represented in the dataset, and closer to the UCD program on the PCA. Japanese and Asian varieties were located at the center of the crescent. In addition, the PCA highlighted the enrichment of our panel in unique 171 accessions not found in the UCD and USDA panel, thus emphasizing its potential to find new phenotypic diversity for fruit quality traits in cultivated strawberry (Fig. 1H, Fig. S5). Heterozygosity decreased in Californian and Floridian cultivars in comparison to European and Asian cultivars. Interestingly, heterozygosity was significantly higher in cultivars and advanced lines of Invenio and in recent European cultivars (released after 1980) (Fig. 1I).

Genetic groups 1, 2 and 3 are colored in green, purple and orange, respectively. Groups 1, 2, 3: Heirloom & related, European mixed group and American & European mixed groups, respectively.

### Fruit quality traits in the diversity panel

A total of 12 fruit quality traits were investigated in the panel of 223 accessions (Table 1). Traits were related to fruit weight (FW); fruit appearance (COL, skin color; UCOL, uniformity of skin color; UFS, uniformity of fruit shape; ACH, position and depth of achenes); firmness (FIRM); composition (TA, titratable acidity; TSS, total soluble solids (Brix units), and BA, the deduced ratio (Brix/TA)); and skin properties (GLOS, glossiness; SR, skin resistance; BRU, bruisedness). Analyses were carried out over two consecutive years, with the exception of the FIRM and SR traits, which were assessed over a single year (Table 1).

**Table 1.** Summary statistics of the 12 fruit quality traits evaluated on the diversity panel in 2020 and 2021. n, number of accessions; CV, Coefficient of Variation; H², broad sense heritability; %GE, percentage of genotype-by-environment interactions in the total variance; r² structure, structuration of the trait as the coefficient of determination of the linear regression between the trait values and the genetic groups. 20-21 indicates the combined values across two years, 2020 and 2021. ns, not significant genotypic effect.

| Trait | Year | n | Range | Mean | $\sigma$ | CV | H<sup>2</sup> | H<sup>2</sup> G1 | H<sup>2</sup> G2 | H<sup>2</sup> G3 | %GE | r<sup>2</sup><br>structure |
| --- | --- | --- | --- | --- | --- | --- | --- | --- | --- | --- | --- | --- |
| Fruit weight (FW, in g) | 2020 | 169 | 1.8 - 30.9 | 12.4 | 5.4 | 43.6 | 0.96 |  |  |  | - | - |
|  | 2021 | 169 | 2.0 - 32.0 | 12.6 | 5.6 | 44.5 | 0.92 |  |  |  | - | - |
|  | 20-21 | 169 | 1.8 - 32.0 | 12.5 | 5.5 | 44.1 | 0.81 | 0.71 | 0.68 | 0.71 | 23.4 | 0.30 |
| Shape uniformity (UFS) | 2020 | 208 | 1 - 5 | 2.7 | 1.3 | 48.0 | - |  |  |  | - | - |
|  | 2021 | 197 | 1 - 5 | 3.1 | 1.3 | 41.5 | - |  |  |  | - | - |
|  | 20-21 | 208 | 1 - 5 | 2.9 | 1.3 | 44.9 | 0.26 | ns | 0.50 | ns | - | 0.05 |
| Achene position (ACH) | 2020 | 209 | 1 - 5 | 3.5 | 0.8 | 23.3 | - |  |  |  | - | - |
|  | 2021 | 198 | 1 - 5 | 3.2 | 0.9 | 28.8 | - |  |  |  | - | - |
|  | 20-21 | 209 | 1 - 5 | 3.4 | 0.9 | 26.2 | 0.66 | 0.71 | 0.67 | 0.60 | - | 0.08 |
| Skin color (COL) | 2020 | 201 | 1 - 7 | 1.2 | 4.507 | 3.7 | - |  |  |  | - | - |
|  | 2021 | 209 | 1 - 7.5 | 1.2 | 4.535 | 3.8 | - |  |  |  | - | - |
|  | 20-21 | 209 | 1 - 7.5 | 1.2 | 4.522 | 3.8 | 0.68 | 0.77 | 0.67 | 0.60 | - | 0.03 |
| Color uniformity (UCOL) | 2020 | 208 | 1 - 5 | 3.0 | 1.3 | 42.7 | - |  |  |  | - | - |
|  | 2021 | 198 | 1 - 5 | 2.8 | 1.3 | 47.0 | - |  |  |  | - | - |
|  | 20-21 | 208 | 1 - 5 | 2.9 | 1.3 | 44.9 | 0.43 | ns | 0.59 | 0.32 | - | 0.10 |
| Firmness (FIRM, in kg/mm) | 2020 | - | - | - | - | - | - |  |  |  | - | - |
|  | 2021 | 210 | 1.04 - 0.07 | 0.433 | 0.2 | 40.3 | 0.98 | 0.94 | 0.97 | 0.95 | - | 0.44 |
|  | 20-21 | - | - | - | - | - | - |  |  |  | - | - |
| Titratable acidity (TA, in g / L) | 2020 | 195 | 0.7 - 2.3 | 1.4 | 0.3 | 20.6 | 0.83 |  |  |  | - | - |
|  | 2021 | 175 | 0.4 - 2.0 | 1.2 | 0.3 | 24.1 | 0.91 |  |  |  | - | - |
|  | 20-21 | 195 | 0.4 - 2.3 | 1.3 | 0.3 | 23.1 | 0.70 | ns | 0.85 | 0.82 | 34.5 | 0.03 |
| Total soluble solids (TSS, in Brix) | 2020 | 207 | 3.6 - 11.0 | 7.1 | 1.4 | 19.7 | 0.92 |  |  |  | - | - |
|  | 2021 | 184 | 3.2 - 11.8 | 6.9 | 1.5 | 22.1 | 0.94 |  |  |  | - | - |
|  | 20-21 | 207 | 3.2 - 11.8 | 7.0 | 1.5 | 20.9 | 0.78 | 0.68 | 0.83 | 0.75 | 30.2 | 0.04 |
| Brix/acidity ratio (BA) | 2020 | 195 | 2.9 - 7.9 | 5.1 | 1.0 | 19.9 | 0.87 |  |  |  | - | - |
|  | 2021 | 175 | 2.1 - 12 | 5.6 | 1.5 | 27.5 | 0.95 |  |  |  | - | - |
|  | 20-21 | 195 | 2.1 - 12 | 5.3 | 1.3 | 24.8 | 0.54 | ns | 0.85 | 0.51 | 48.5 | 0.00 |
| Glossiness (GLOS) | 2020 | 195 | 1 - 5 | 3.5 | 1.2 | 35.8 | - |  |  |  | - | - |
|  | 2021 | 182 | 1 - 5 | 3.5 | 1.1 | 31.8 | - |  |  |  | - | - |
|  | 20-21 | 195 | 1 - 5 | 3.5 | 1.2 | 33.8 | 0.77 | 0.76 | 0.64 | 0.43 | - | 0.48 |
| Skin resistance (SR) | 2020 | - | - | - | - | - | - |  |  |  | - | - |
|  | 2021 | 210 | 0 - 3 | 1.65 | 1.0 | 58.2 | - |  |  |  | - | 0.34 |
|  | 20-21 | - | - | - | - | - | - |  |  |  | - | - |
| Bruisiness (BRU) | 2020 | 199 | 1 - 5 | 2.5 | 1.3 | 50.8 | - |  |  |  | - | - |
|  | 2021 | 54 | 0 - 5 | 1.8 | 1.0 | 54.3 | - |  |  |  | - | - |
|  | 20-21 | 199 | 0 - 5 | 1.9 | 1.1 | 55.7 | 0.57 | 0.52 | 0.45 | ns | - | 0.49 |

Most of the traits exhibited a considerable range of variation in the diversity panel, with coefficients of variation ranging from 3.7% for COL to 58.2% for skin resistance. For example, FW (average: 12.5 g) ranged from 1.8 and 32 g. (Table 1). Most traits showed a normal distribution, while SR, GLOS, UFS and BRU showed a skewed distribution (Fig. S6). Nine traits displayed high amount of genotypic variance associated with high broad sense heritability (H^2^) ranging from 0.66 (ACH) to 0.98 (FIRM); H^2^ of four traits, namely UFS (0.26), UCOL (0.43), BA (0.54) and BRU (0.57), was below 0.6 (Table 1). Few variations of H^2^ between groups were observed for FW and FIRM, suggesting that phenotypic variability was equivalent between groups, whereas a strongest decrease in H^2^ was observed for GLOS and BRU in G3 (Table 1). A significant interaction between genotype and environment was detected for all the traits for which repeated measurements were available over two years (Table 1), with the effect of environment being strongest for traits related to fruit composition (TSS, TA, BA).

To further explore the phenotypes of the diversity panel, we performed a PCA of the 223 accessions using a PCA biplot (Fig. 2A). PC scores revealed that the three genetic groups were distributed differently according to PC1 (39.2%) and PC2 (17.2%) in terms of fruit quality traits. G1 was distinct from G2 and G3 (Fig. 1B). Examination of the loadings of the traits on PC1 further showed that FW, appearance (UFS, UCOL), FIRM and skin properties (GLOS, SR and BRU) traits were responsible for the separation between G1 on one side and G2 and G3 on the other side. TSS, TA and ACH had a very small contribution to the differentiation of the three subpopulations along PC1, and mostly contributed to PC2 and PC3, respectively (Fig. 2A, Fig. S7).

**Figure 2.**
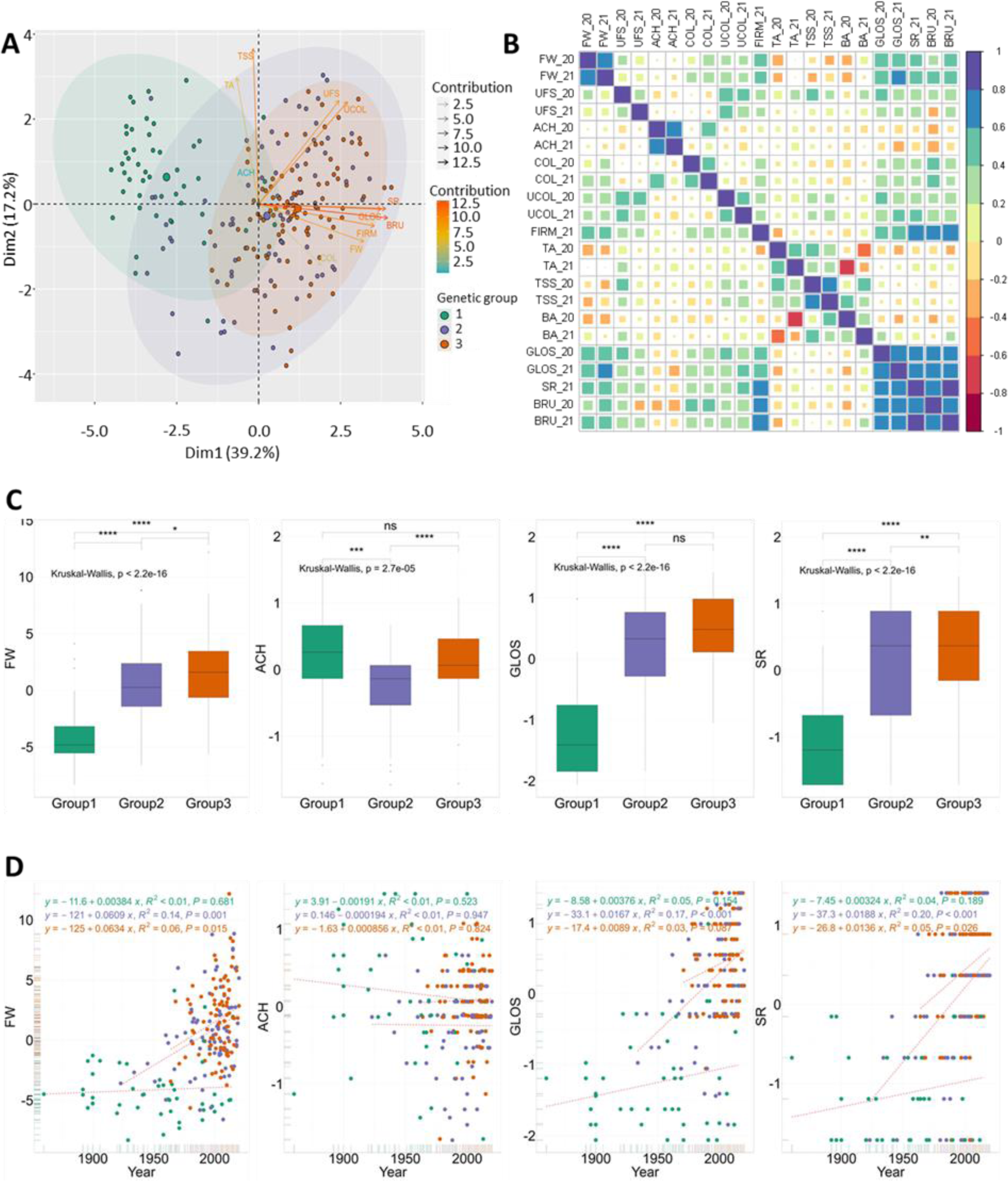
Phenotypic variations across the three genetic groups of the panel. (A) Principal Component Analysis of the 2-year BLUP values for 11 traits. (B) Correlations between the 12 traits for each year. (C) Comparisons of 2-year BLUP values for FW, ACH, GLOS and SR among genetic groups. (D) Genetic gains for FW, ACH, GLOS and SR among genetic groups. Genetic groups 1, 2 and 3 are colored in green, purple and orange, respectively. Groups 1, 2, 3: Heirloom & related, European mixed group and American & European mixed groups, respectively. FW, fruit weight; UFS, uniformity of fruit shape; COL, skin color; UCOL, uniformity of skin color; ACH, position and depth of achenes; FIRM, firmness; TA, titratable acidity; TSS, total soluble solids; GLOS, glossiness; SR, skin resistance; BRU, bruisedness.

Correlation analysis of fruit quality trait data collected over 2020 and 2021 (Fig. 2B, Table S2) supported the relationships identified in the PCA biplot (Fig. 2A). GLOS, SR and BRU traits were positively and strongly correlated with each other and with FW and FIRM (*r* = 0.51 to 0.87), indicating the strong potential for directional selection of these traits (Fig. 2A). UFS and UCOL were also highly correlated among them (*r* = 0.64) and, to a lesser extent (*r* = 0.35 and 0.26), with FW. TSS and TA were significantly correlated (*r* = 0.45) but, within groups, the correlation was only significant for G2 (*r* = 0.51) and G3 (*r* = 0.47) groups (Fig. S8). FW also demonstrated significant negative correlations with TSS (*r* = −0.21) and TA (*r* = −0.15) (Fig. 2B; Table S2). No or weak correlations were observed between ACH and other fruit quality traits.

Most fruit quality traits have undergone significant phenotypic changes over time, as old varieties have evolved into modern cultivars (Figs 2C, 2D, Fig. S9). Phenotypic values of all fruit quality traits, except BA and COL, were significantly different between the three genetic groups. For example, FW considerably increased during the modern breeding phase, as reflected mainly in trends within G2 and G3 (Figs 2C, 2D) G1 was associated with low FW, dull, soft, low SR and easily wounded skin with uneven color and shape, whereas G3 exhibited the highest values for these traits (Fig. 2C, Fig. S9). G2 was equivalent to G3 for UFS and UCOL, TSS and TA, and GLOS; and was in the average of G1 and G3 for FW, FIRM, SR and BRU. Cultivars from G2 displayed more outcropped achenes than the others (Fig. 2C, Fig. S9).

These changes are linked to significant genetic gains over time for most fruit quality traits, with the exception of ACH and BA (Fig. 2D, Fig. S10). However, while positive and time-dependent genetic gains were observed within G2 and G3, genetic gains were usually low or inexistent within G1. For FW, for example, several recent varieties of the G1 showed the same trait values as old ones. A negative, non-significant trend was even observed for TSS and TA, whose values were lower in G2 and G3 than in G1 (Fig. S9). Remarkably, regardless of the TSS and TA reduction in modern varieties compared to old varieties, no significant differences for BA values were observed between groups (Fig. S9).

### GWAS of fruit quality traits

To reveal the genetic architecture of fruit quality in strawberry, we performed GWAS on the 12 fruit quality traits assessed in the 223 accessions of the strawberry diversity panel using genome-wide SNP markers from the 50K FanaSNP array^21^. The structuration of the population (Figs 2B, 2C) was considered by fitting both kinship and structure as cofactors for GWAS analysis. Detailed Manhattan plots for all 12 traits are shown in Figs 3, 4, Fig. S11. The 71 significant associations with SNP markers are distributed on 51 chromosome regions spread on 23 chromosomes (Table S3).

### Fruit weight and appearance (FW, UFS, COL, UCOL, ACH)

Three significant SNP were identified for FW on chromosome 1B (19,119,571 bp, p-value 6.74E-09), 5B (17,045,086 bp, p-value 1.60E-06) and a highly significant SNP on chromosome 2D (15,565,564 bp, p-value 3.27E-12) (Fig. 3A, Table S3). The minor allele of AX-184592155 had a phenotypic variance explained (PVE) of 11.8% with an effect of 1.8g on FW (Fig. 3B, Table S3). Sixteen unique significant SNPs were identified for appearance traits, eight for UFS, three for ACH and five for COL (Fig. 3A, Fig. S11, Table S3). No signal was detected for UCOL.

**Figure 3.**
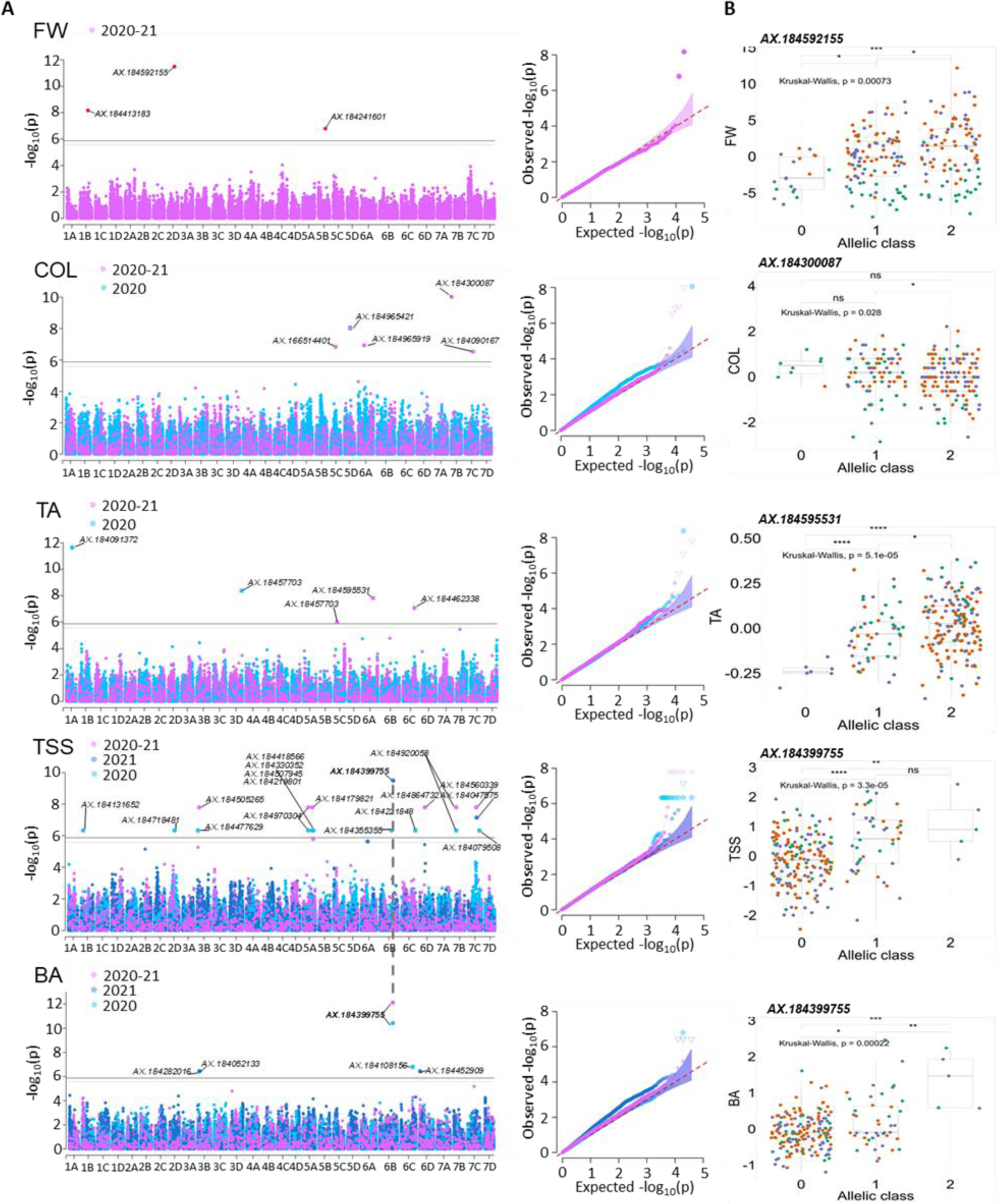
Genome wide association study of FW, COL, TA, TSS and BA. (A) Manhattan and Q-Q plots for yearly and 2-year BLUP values. (B) Effect of the most significant SNP marker. Genetic groups 1, 2 and 3 are colored in green, purple and orange, respectively. Groups 1, 2, 3: Heirloom & related, European mixed group and American & European mixed groups, respectively. FW, fruit weight; COL, skin color; TA, titratable acidity; TSS, total soluble solids; BA, brix acidity ratio. Marker classes are as follows: 0=AA genotype, 1=AB, and 2=BB genotype according to the Axiom™ Strawberry FanaSNP 50k.

#### Fruit composition (TA, TSS, BA)

Twenty-seven unique significant SNPs were detected for fruit composition traits, five for TA, 18 for TSS and five for BA (Fig. 3A, Table S3). The SNP AX-184595531 detected for TA on the 2020-2021 combined values (25,621,066 bp, p-value 1.55E-08) was particularly notable for its PVE of 35.5%. Only seven cultivars, all belonging to G2, were unfavorable homozygous for this marker (Fig. 3B). SNPs AX-184091372 on chromosome 1A (13,540,517 bp, p-value 2.30E-12, PVE 17.1%) and AX-184457703 on chromosome 3D (25,621,066 bp, p-value 4.22E-09, PVE 17.9%) were also of particular interest for their PVE and impacting effect on TA in 2021. AX-184399755 was the highest effect SNP for TSS in 2021 on chromosome 6B (31,578,303 bp, p-value 3.24E-10, PVE 19%) (Fig. 3B). It was also highly significant for BA in 2020 (p-value 3.84E-11, PVE 46.1%) and 2020-2021 combined values (p-value 7.88E-13, PVE 65.8%) (Fig. 3B). Only five cultivars were favorable homozygous for this marker.

Interestingly, on diploid *F. vesca* reference genome (*F. vesca* v4.0.a1^32^), the position of SNPs AX-184399755 (6B) (10,248,453 bp on diploid) was close (606,797 bp apart) to the position of AX-184864732 (6D) (10,855,248 bp on diploid) indicating that these two TSS QTL could be homoeo-QTL. Same observation was made for SNP AX-184920058 (7B) (20,136,538bp on diploid) and AX-184079508 (7C) (19,687,318 bp on diploid), both being associated to TSS QTL.

#### Fruit firmness and skin properties (FIRM, GLOS, SR, BRU)

Four significant SNPs were detected for FIRM, six for GLOS, five for SR and five for BRU (Fig. 4A, Table S3). The chromosome 3D was of particular interest for these traits as it comprises one highly significant SNP for FIRM (29,275,014 bp, p-value 6.07E-12, PVE 11.2%) and the highly significant SNP AX-184177060 (27,845,440 bp) common to both GLOS (p-value 9.01E-10, PVE 26.2% on combined values) and SR (p-value 6.45E-07, PVE 8.4%) (Fig. 4B). The latter SNP was detected systematically in 2020, 2021 and 2020-2021 for GLOS with PVE ranging from 26.2% to 28.7%, with a negative effect of the minor allele ( −0.7 to −0.8 on 1 to 5 scale). MAF of this SNP was highly reduced towards G3, indicating strong selection of the favorable allele (Fig. 4B). SNPs for BRU bruisedness were detected for the 2020-2021 combined values only.

**Figure 4.**
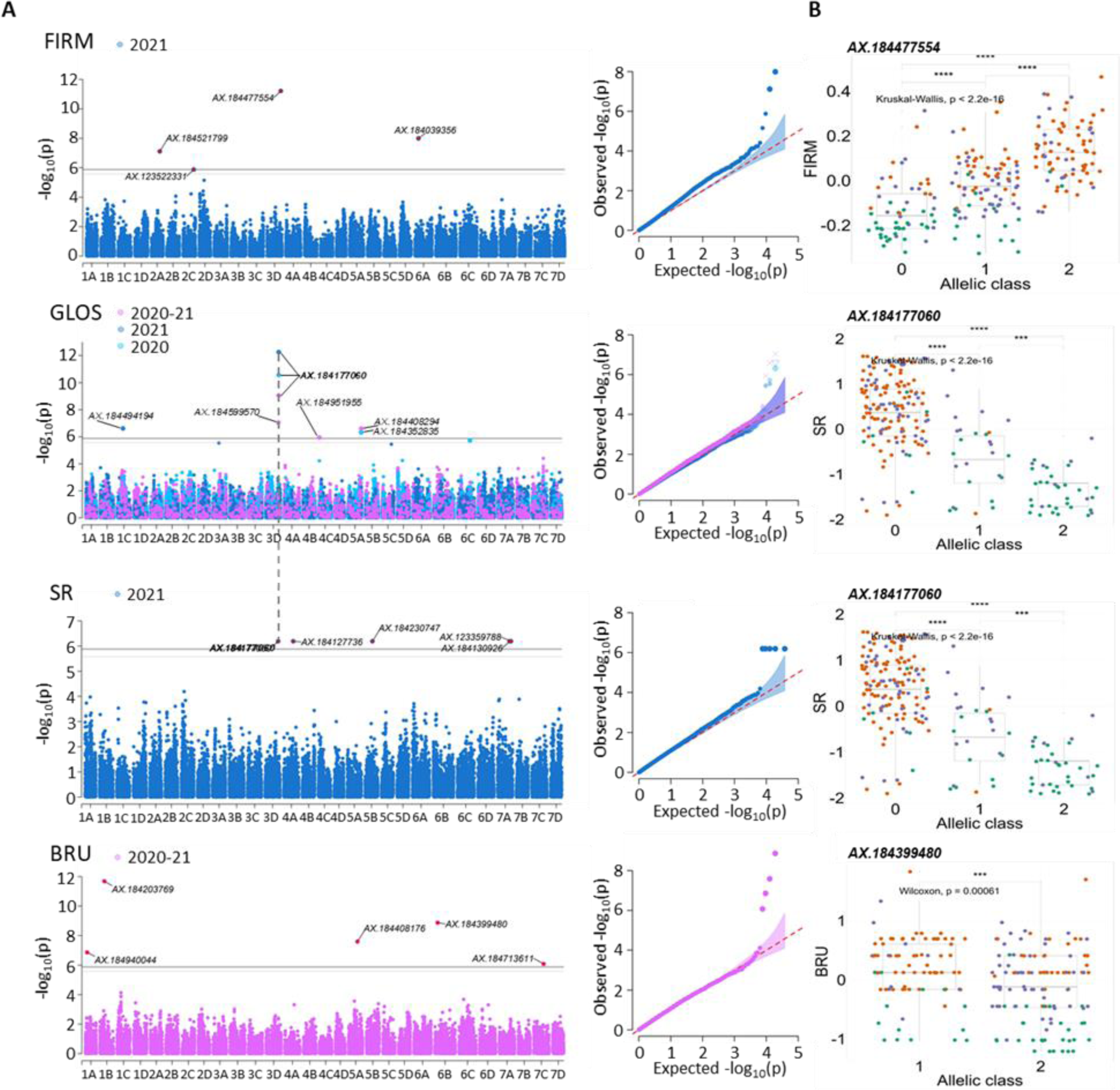
Genome wide association study of FIRM, GLOS, SR and BRU. (A) Manhattan and Q-Q plots for yearly and 2-year BLUP values. (B) Effect of the most significant SNP marker. Genetic groups 1, 2 and 3 are colored in green, purple and orange, respectively. Groups 1, 2, 3: Heirloom & related, European mixed group and American & European mixed groups, respectively. FIRM, firmness; GLOS, glossiness; SR, skin resistance; BRU, bruisedness. Marker classes are as follows: 0=AA genotype, 1=AB, and 2=BB genotype according to the Axiom™ Strawberry FanaSNP 50k.

### Selective sweep signals during strawberry improvement

We identified markers under selection during strawberry improvement in light of genome scans based on Mahalanobis distance on the whole diversity panel (Fig. 5A) and nucleotide diversity among genetic groups throughout the genome (Fig. 1F, Fig. S3). Six associations were found among candidate selective sweeps based on Mahalanobis distance (Fig. 5A, Table S4), one for UFS, one for FIRM, one for TSS, two for GLOS and one for SR, where SNPs AX-184177060 associated to GLOS and SR, AX-184477554 associated to FIRM, and AX-184864732 associated to TSS, presented a drastic reduction in nucleotide diversity towards modern genotypes (Fig. 1F, Fig. S3, Table S4). For example, in the case of the SR and GLOS SNP marker AX-184177060, the favorable allele is over-represented in the most recent accessions (average year of release 2000), whereas cultivars heterozygous and unfavorably homozygous for the marker were respectively released on average in 1967 and 1947, supporting the fact that the favorable allele has been selected over time. Seven others markers associated to UFS, TA, TSS, GLOS and BRU were also found in genomic regions with substantial depletion in nucleotide diversity in G3 compared to the other groups.

**Figure 5.**
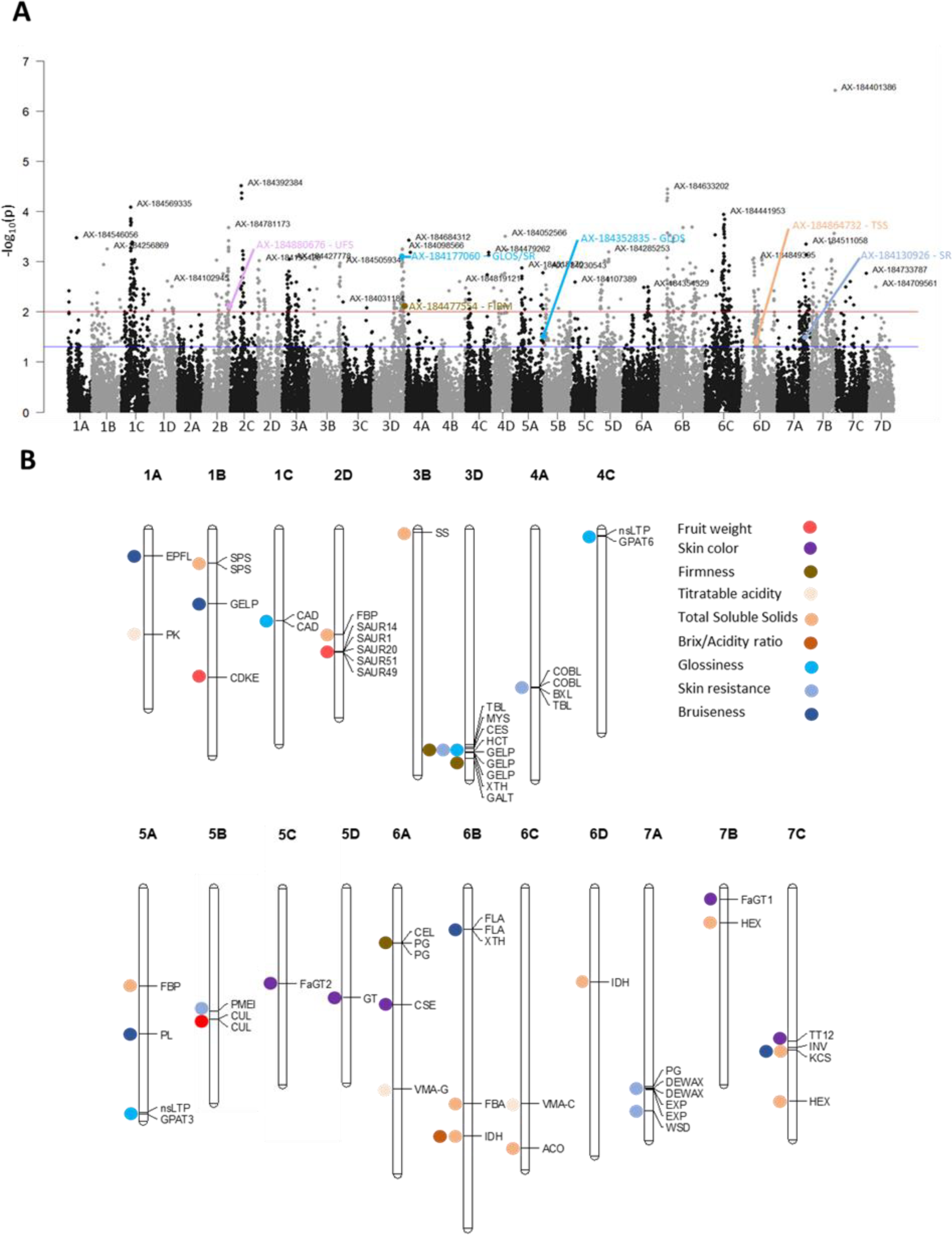
Selective sweeps and candidate genes. (A) Selective sweeps as a Manhattan plot of p-values of the genome scan based on Mahalanobis distance. Red and blue lines indicate thresholds at 0.01 and 0.05, respectively; trait associations with a p-value < 0.05 are indicated with colors. (B) Physical mapping of the candidate genes underlying the GWAS of nine traits on the Camarosa genome.

### Candidate genes were identified for 37 QTL controlling 9 fruit quality traits

Candidate genes (CG) underlying fruit quality QTL were identified within a window of ∼400 kb surrounding the QTL marker. This value, which corresponds to the short-range LD found in the Californian *F.* × *ananassa*^4^, is stringent compared to the average LD calculated on the 28 linkage-groups in our diversity panel, which is 932 kb. In chromosome regions harbouring strong QTL of interest and displaying low genetic diversity and high LD, i.e. the 3D region extending from 23,233 to 29,635 kb (Fig. 1F), we considered much larger intervals based on LD estimates (up to ∼1,382 kb) for 3B and 3D. We excluded two traits (UFS and ACH) from CG analysis because the molecular pathways underlying these traits are far from being deciphered in strawberry. No QTL was detected for UCOL. In total, we identified 64 candidate loci for 37 SNP markers associated with the nine fruit quality traits (Fig. 5B). Table 2 provides names, abbreviations and positions on Camarosa and Royal Royce genomes of these 64 CG. Their possible functions are indicated in Table S5.

**Table 2.**
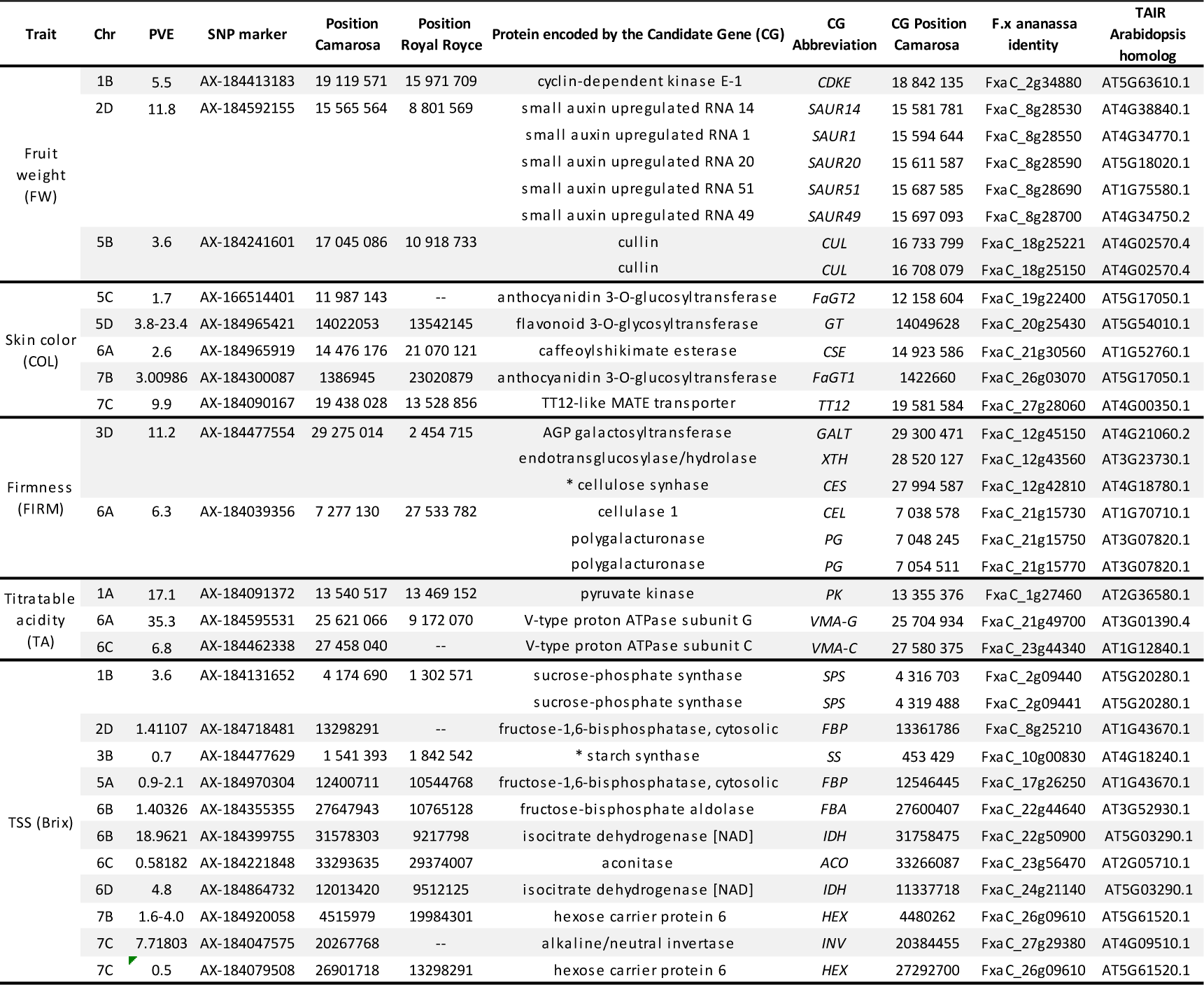
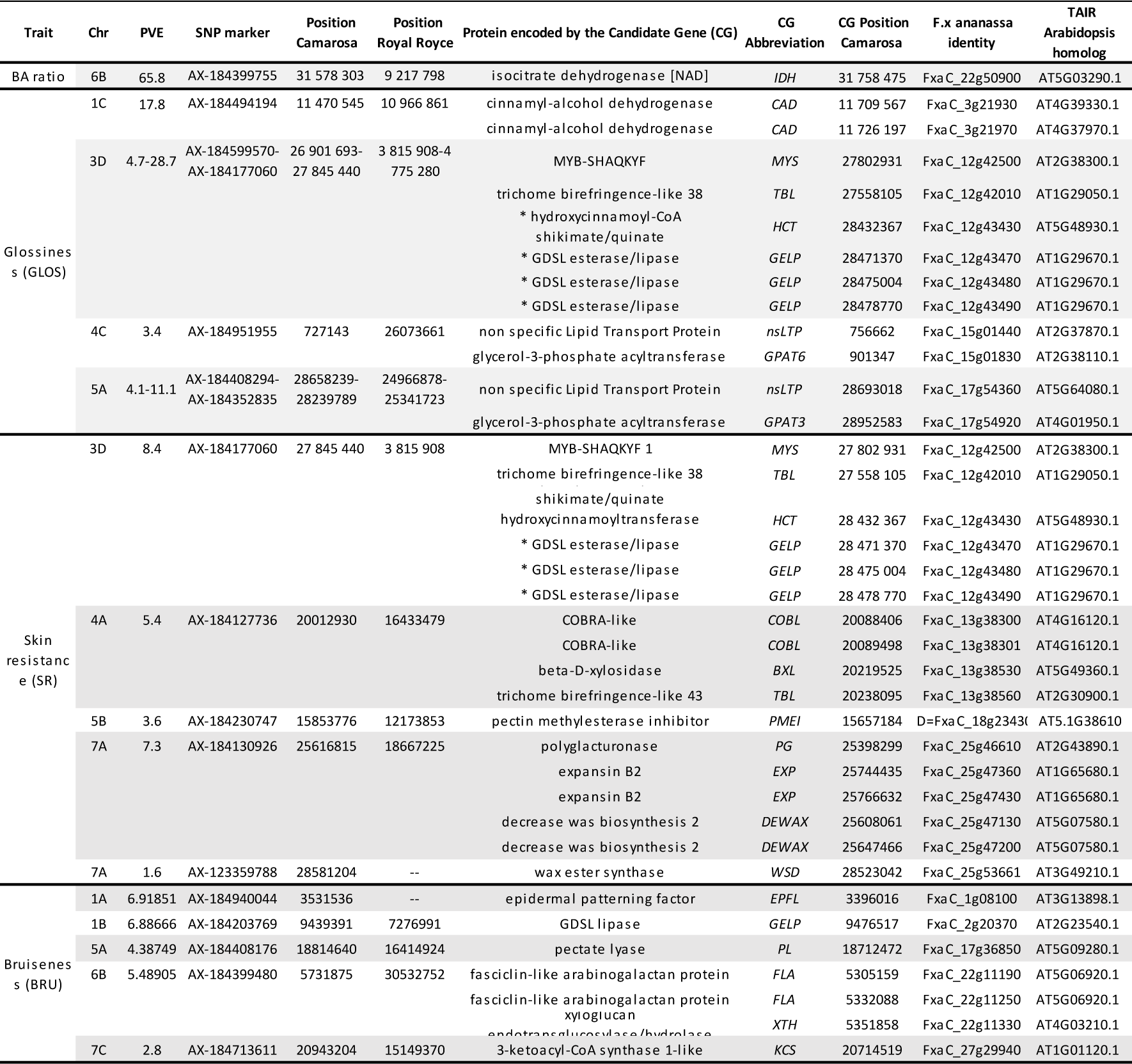
Candidate genes underlying nine fruit quality traits. PVE, Phenotypic variance explained (%). Position Camarosa and Position Royal Royce: physical positions on Camarosa and Royal Royce reference genomes. Protein encoded by the candidate gene (CG): annotation in the Camarosa genome. CG Position Camarosa and *F.x ananassa* identity: physical position and identity in the Camarosa genome. * indicate the CG that were found outside the ∼400 kb interval around the SNP marker.

## Discussion

### Genetic and phenotypic shifts in modern strawberry breeding programs

Our study sheds light on the genetic and phenotypic shifts that occurred over the last 160 years of strawberry breeding by analyzing 223 accessions comprising original old and modern European breeding material. According to our analyses, old strawberry cultivars, which here consist mainly of European cultivars selected before 1950 and included in the Heirloom & related group (G1), are clearly separated from other genetic resources (Fig. 1C), in agreement with earlier studies^31^ confirmed in recent papers^4,33^. For over half a century^34^, breeding programs in Western and Southern Europe have made extensive use of Californian cultivars and, more recently, of Floridan cultivars, which are underrepresented in our study, as progenitors. As a consequence, our results show the clustering of most European recent cultivars in an American & European mixed group (G3). The European mixed group (G2), which includes other European cultivars, is likely related to the group previously named Cosmopolitan^4^. However, European from US subpopulations were separated in the USDA panel study, thanks to the large number of US accessions^27^.

The nucleotide diversity is overall well conserved among the genetic groups of our panel (Fig. 1E). In contrast, a significant erosion of genetic diversity was observed in highly structured populations^4,5,28^. We found a more nuanced picture by examining nucleotide diversity at the chromosome level, since it drops dramatically in regions potentially subject to selection pressure (Fig. 1F, Fig. S3). The diminution in both LD and heterozygosity specifically observed in the most recent American cultivars^4^ is likely explained by the gradual closure of Californian and Floridian programs. In contrast, cultivars and advanced lines of Invenio as well as the recent European cultivars released after 1980 display higher heterozygosity values (Fig. 1I). One possible explanation for the high genetic diversity retained in European accessions is that European breeders had to cope with a wide range of breeding targets due to the diversity of cultural practices, markets and consumer preferences found in Europe^6,35^. High quality strawberry varieties released in Europe therefore had to meet the requirements of both high cultivar performance, e.g. high fruit yield, as in Californian cultivars^5^ and high sensory fruit quality, e.g. high flavor^6^.

Remarkably, recent studies have shown that despite a loss in genetic diversity, increases in both genetic gain and phenotypic variation were observed in highly structured populations such as those of the Californian breeding programs^5^. In these programs, breeding efforts rapidly led to the improvement of fruit weight and fruit firmness^5,26,28^, which participated to the so-called Californian green revolution^5^. European breeding programs have benefited from these efforts, as modern American cultivars appear in the pedigree of prominent European cultivars^3^. Consistently, our results indicate a similar trend towards improved fruit size and firmness, as well as skin glossiness and resistance, in recent European germplasm (Fig. 2C, Fig. S9). Interestingly, we found that TSS and TA values decreased over time in G1 but that the BA ratio kept the same value (Figs S9, S10), in agreement with ^5^ who even observed an increase in BA levels, which could partly counterbalance the decrease in fruit sweetness. According to ^5,26,30^, this reflects the antagonism between yield and firmness, on one side, and TSS and TA, on the other side.

### Novel markers for the selection of fruit quality traits

GWAS is a powerful tool for the detection of SNP markers linked to different traits in strawberry^9,17,28,30,36,37^. Here, based on a large diversity panel, we detected 71 markers associations to major fruit quality breeding targets. Some of the marker/QTL associations detected confirm published results and, consequently, validate our findings in a different genetic context. For example, our AX-184039356 marker linked to FIRM on 6A is very close to those previously described for fruit firmness^4,17^. Likewise, our AX-184477629 marker linked to TSS on 3B is in the same chromosome region as the SSC1 QTL controlling soluble solids content^30^. However, we found in GWAS/QTL published data only a few additional fruit quality QTL located on the same chromosomes as those detected here. Among them are the FW QTL on 5B and the TSS QTL on 5A previously reported^38^ on the same chromosomes but in different regions. Thus, thanks to the originality of European accessions included in our study, the genetic diversity of the panel allowed us to reveal new QTL and associated SNP markers, even for well-studied fruit quality traits such as FW and TSS.

In contrast to these well-studied traits, few studies have unveiled the genetic architecture of skin associated traits such as fruit glossiness^17,39^ which is, alongside color, one of the most prominent traits for fruit attractivity to the consumer^40^. Remarkably, among the six associations found among the selective sweeps detected, two were found for GLOS and one for SR (Fig. 5A). Furthermore, by highlighting a ∼6,400 kb region on chromosome 3D linked to glossiness, skin resistance and firmness, our results shed a new light on a genomic region under strong breeding pressure (Figs 1F, 5A). Remarkably, among the six associations found among candidate selective sweeps, two were found for GLOS and one for SR. This chromosome region has thus probably played a crucial role in improving the attractivity and post-harvest qualities of strawberries, a feature that is receiving increasing attention in strawberry breeding programs. Information on the position of SNP markers on both Camarosa and Royal Royce genomes will facilitate new studies on fruit quality traits, thus contributing to validate these markers for MAS.

### Candidate genes

#### Fruit weight and appearance

Fruit weight (FW) and shape are complex traits. Underlying genes of previously unknown functions have been identified by map-based cloning in species such as tomato^41^ and corresponding CG have been detected in several crops^42^. Translation of these findings to strawberry may however prove difficult because of the different ontogenic origin of strawberry, which is an accessory fruit derived from the flower receptacle and not from the ovary. Indeed, our GWAS study did not detect any known gene families linked to fruit weight and shape, but highlighted for FW QTL several CG (*CDKE,* a cluster of five *SAUR*, *CUL*) involved in cell division and expansion processes and their regulation (Table S5).

Red-colored anthocyanins, which give strawberries their attractive bright red appearance, are flavonoids derived from the phenylpropanoid pathway. In cultivated strawberry, allelic variants of the master regulator *MYB10* belonging to the MBW complex have been shown to be responsible for the white skin-color and red flesh-color^8,11^. In our GWAS study, we did not detect any previously known color QTL nor CG linked to the MBW complex, probably because white fruit genotypes and flesh-color trait were under-represented in our analysis. However, our diversity panel has enabled us to reveal new skin color QTLs and identify strong CG involved in the successive steps leading to anthocyanin accumulation in strawberry^11^: (i) anthocyanin biosynthesis; (ii) formation of stabilized anthocyanidin-glucosides; and (iii) transport of anthocyanidin-glucosides for storage in the vacuole. Color CG, which deserve further study, include a gene (*CSE*) encoding shikimate esterase, an enzyme involved in lignin pathway that may compete with anthocyanin biosynthesis for common substrates; several genes encoding glycosyltransferases (*GT*), among which the strawberry FaGT1 enzyme that has been shown to generate anthocyanidin 3-O-glucosides^43^ and its homolog FaGT2; and a gene encoding a vacuolar flavonoid/H+-antiporter (*TT12*) that can actively transport cyanidin-3-O-glucoside to the vacuole^44^.

#### Fruit firmness and composition

Breakdown of the cell wall (CW) is the main mechanism responsible for fruit softening during ripening. CW is mainly constituted by a cellulose-hemicellulose network immersed in a pectin matrix. Strawberry fruit softening involves the pectin-degrading enzymes polygalacturonase (PG) and pectate lyase (PL)^45^. The down-regulation of *PL*^46^ and of *PG*^47^ influences fruit firmness and/or shelf life of strawberry. Many additional proteins are involved in CW modifications e.g. pectin methylesterase (PME) and its inhibitors (PMEI) that control cell adhesion and elasticity through pectin esterification, enzymes of the xyloglucan endotransglycosylase/hydrolase (XTH) family involved in hemicellulose remodelling, cellulases (CEL) that degrade cellulose, and expansins (EXP) that promote CW loosening. Other enzymes such as cellulose synthase (CES) or proteins with ill-defined roles such as arabinogalactan-proteins (AGPs) likely play a role in CW structure and properties. Therefore, considerable variations in fruit firmness can be expected by modulating the activity of enzymes encoded by CG underlying the 3D QTL (*GALT, XTH, CES*) and 6A (*CEL, PG*) QTL. *XTH* and *CES* are strong candidates located at 750 to 1280 kb from the AX-184477554 marker in the well-conserved 3D region while *CEL* and *PG* underly the 6D FIRM QTL previously detected^4^.

The sugar/acid balance is central for consumers perception of fruit quality^19^ and the sugar/acid ratio has been widely adopted as a breeding target^5^. The major soluble sugars that accumulate during fruit ripening are glucose, fructose and sucrose, the concentration of which depends on the cultivar^48^. The major organic acids are malate and especially citrate, which is the predominant organic acid^48^. Their concentrations are stable or decrease during fruit ripening. Fruit sweetness is usually assessed in refractometer (Brix units), which measures total soluble solids (TSS), including sugars and organic acids. Fruit acidity is assessed by titratable acidity (TA), to which citrate contributes most in strawberry. The accumulation in strawberry of soluble sugars and organic acids depends on synthesis in the leaf (source) and long-distance transport of photoassimilates (sucrose, inositol) to the fruit (sink). Photosynthetic sugars are further metabolized in the fruit to produce soluble sugars and organic acids that are then stored in the vacuoles^49^. Our GWAS study identified several CG implicated in the metabolism of sugars, either in the leaves or in the fruit, including *SPS* (1B QTL), *FBP* (2D and 5A QTL), *SS* (3B QTL), *FBA* (6B QTL), and *INV* (7C QTL). The starch synthase (*SS)* is located more than 1 Mb apart from the 3B QTL marker but has been recently identified as a CG for a TSS QTL^30^. The neutral invertase (*INV*), which underlies the major 7C TSS QTL (PVE 7.7%), is a strong candidate that has been shown to be crucial for glucose and fructose accumulation during ripening in tomato^50^ while a cell wall invertase is responsible for a major TSS QTL in this species^51^. Another strong candidate is the hexose transporter (*HEX*) (7B and 7C QTL), which could transport glucose and fructose across the tonoplast, as suggested in grape berries^52^. Furthermore, the *HEX* gene may underlie two possible TSS homoeo-QTL located on chromosomes 7B and 7C, respectively.

Two CG underlying TSS QTL encode enzymes involved in the tricarboxylic acid (TCA) cycle, notably isocitrate dehydrogenase [NAD] (IDH) (6B QTL) and aconitase (ACO) (6C QTL). TCA is the central metabolic cycle that uses substrates from the glycolysis to produce energy. It fulfills major roles in the fruit, among which the metabolism of citric acid^53^. While aconitase has been shown to contribute to the regulation of acidity in the citrate-accumulating lemon^54^, we did not detect any TA QTL corresponding to the 6C Brix QTL. Interestingly, *IDH* underlies strong shared QTL for TSS (PVE=19.0) and BA (PVE=65.8) on chromosome 6B. The implication of IDH a significant contributor to the TCA cycle, in the sugar/acid balance of strawberry, therefore merits further studies. Moreover, *IDH* is also located at ∼ 675 kb from the TSS QTL on chromosome 6D, indicating that it could underlie two TSS homoeo-QTL located on chromosome 6B and 6D, respectively. As previously suggested^13^, the detection of homoeo-QTL could depend on environmental conditions, which vary according to the year of study.

The CG underlying the 1A TA QTL encodes pyruvate kinase (PK) a crucial enzyme for gluconeogenesis which has already been demonstrated to regulate citric acid metabolism during strawberry fruit ripening^53^. Two additional CG for the TA QTL located on 6A (PVE 35.3%) and 6C (PVE 6.8%) encode subunits of the V-type proton ATPase (*VMA-G* and *VMA-C*), respectively. Both are strong candidates for the control of fruit acidity, as they are part of a protein complex whose role is to generate a proton gradient across the tonoplast, which is essential to drive the storage of organic acids in the vacuole of fleshy fruits^55^.

#### Skin properties

The outermost wall of the fruit is composed of the cuticle, the epidermis and several layers of sub-epidermal cells^56^. This ill-defined tissue, also called fruit skin^57^, acts as a barrier against water-loss and pathogens and provides protection against mechanical injuries^58^. Its properties depend on epidermal and sub-epidermal cell patterning (cell size and shape) and on the composition and structure of CW and cuticle. To date, the cuticle has been poorly studied in strawberry, except for its composition^59^. Recent studies, in particular in the tomato model, furthered our understanding of the synthesis of cuticle components (wax and cutin polyester, phenolics) and explored the complex interactions between cutin polyesters, CW polysaccharides and phenolics, and their possible contribution to cuticle properties^56^.

Fruit glossiness is an environment-sensitive trait linked to wax and cutin accumulation on the fruit surface but also to epidermal cell patterning^60^. Among CG identified for GLOS QTL are genes involved in phenylpropanoid pathways (*CAD* in 1C QTL), epidermal patterning (*TBL* in 3D QTL), regulation of wax biosynthesis (MYS, 3D QTL), lipid and cutin biosynthesis (GPAT6, 4C QTL; GPAT3, 5A QTL) and possibly transport of cutin precursors (nsLTP, 4C and 5A)^58,61^. In addition to the *MYS* gene, a transcription factor involved through *DEWAX* in the regulation of the ECERIFERUM1 (CER1) enzyme involved in the biosynthesis of wax alkanes^62^, this region harbors, within ∼700 kb of 3D QTL markers, the phenolic pathway *HCT* gene that is essential for cuticle formation^63^ and, close-by, three *GELP* genes. Several members of the large *GELP* family have been demonstrated to play crucial roles in cutin polymerization (cutin synthase^64^) and in assembly-disassembly of the related polyester suberin^65^. Examination at the Tomato eFP Browser (http://bar.utoronto.ca) and TEA-SGN (https://tea.solgenomics.net) databases of the expression of the three tomato closest homologs (*Solyc03g005900, Solyc02g071610, Solyc02g071620*) of the 3D GLOS QTL-linked *GELP* genes indicate that they are strongly expressed in the young fruit, when the cutin synthesis rate is the highest^60^. Furthermore, in the tomato pericarp, their expression is restricted to the outer and inner epidermis. These findings strongly suggest that, in cultivated strawberry, a cluster of genes with likely roles in cuticle formation and structure has been selected in modern varieties for its impact on fruit cuticle-related traits, including GLOS.

Remarkably, we found that the major skin resistance (SR) QTL, which estimates the fragility of the fruit surface to peel off when a mechanical stress is applied, is shared with the GLOS QTL on 3D. The major 3D FIRM QTL (PVE 11.2%) was also found nearby (at ∼1400 kb). Since the FIRM trait was estimated by measuring the force needed to punch a hole in the fruit surface (penetrometer), it can be linked to the properties of the fruit skin. Interestingly, connections between fruit firmness and the cuticle have recently been demonstrated in tomato where changes in cuticle composition and properties are responsible for a major firmness QTL^66^. Altogether, these results suggest that in the 3D conserved region, modifications of fruit surface properties, either due to changes in epidermal cell patterning and/or in cell wall and cuticle properties, have been selected in modern strawberry varieties for their effect on both fruit glossiness, resistance to mechanical damages, and possibly firmness. Other candidates linked to either epidermis patterning (*TBL* on 4A), cell wall modifications (*COBL* and *BXL* on 4A, *PMEI* on 5C, *PG* and *EXP* on 7A) and cuticle formation (*DEWAX*, a target of MYS, and *WSD* on 7A) underly the additional SR QTL detected.

In contrast, none of the QTL detected for fruit bruisedness (BRU), a trait assessed visually, were found to co-localize with either GLOS, SR or FIRM QTL while all these traits are strongly correlated, indicating that the underlying mechanisms are probably different or that the corresponding QTL are below the detection threshold. CW-related GC that may affect cell wall properties (*PL* on 5A, *XTH* on 6B) or cell adhesion of sub-epidermal cells (*FLA* on 6B^67^) merit further investigation, as fruit susceptibility to bruising is essential for post-harvest handling and defense against fruit decay.

## Conclusion

In summary, the exploration of untapped genetic resources, including original European cultivars spanning 160 years of breeding, has revealed considerable changes in recent decades in the genetic and phenotypic diversity of cultivated strawberry. American cultivars have had a major impact on recent European breeding programs and, therefore, on modern strawberry varieties in Europe. However, our findings also revealed that a considerable, and previously undescribed, genetic diversity can be harnessed for improving fruit quality through breeding. While most fruit quality traits are involved, our study underlines the contribution of little-studied traits related to the fruit surface (glossiness, skin resistance, bruisedness) to the breeding of modern varieties. For these and other traits, the strong CGs underlying the main QTL detected warrant further investigation, for example through additional association studies or functional analyses. From a more applied perspective, the genetic markers highlighted will be used for the selection of improved strawberry varieties with high fruit quality.

## Materials and methods

### Plant materials and experimental design

A total of 223 accessions from the historical germplasm collection of Invenio was chosen to constitute the diversity panel. The trial took place in a soilless system, at Douville in the South-West of France (45° 1.2831’ N; 0° 37.0198’ E, France). The crop management was the one used for commercial semi-early cultivated strawberry in France. The trial was organized in a randomized complete block design of two blocks of four biological replicates each in a 288 m² glass greenhouse in 2020 and 2021. Planting of tray plants occurred around the 15^th^ of December of the previous year.

### Sample preparation and phenotyping

Fruits were harvested once per season and evaluated for 12 fruit quality traits: FW, fruit weight; UFS, uniformity of fruit shape; COL, skin color; UCOL, uniformity of skin color; ACH, position of achenes; FIRM, firmness; TA, titratable acidity; TSS, total soluble solids; BA, TSS/TA ratio; GLOS, glossiness; SR, skin resistance;

BRU, bruisedness. FW was evaluated as the mean weight of harvested fruits after discarding immature and overripe fruits. UFS, UCOL, ACH as well as GLOS and BRU were visually assessed on 1-5 scales (Table 1) as a single note on a whole strawberry tray (>10 red ripe fruits). COL was evaluated on 4-5 red ripe fruits a 1-8 scale based on the strawberry color chart from Ctifl (http://www.ctifl.fr/Pages/Kiosque/DetailsOuvrage.aspx?IdType=3&idouvrage=833). FIRM was evaluated on six fruits from each accession with an FTA-GS15 (Güss) penetrometer (5 mm diameter) at 3 mm depth (5 mm/s speed, 0.06kg release threshold). SR was evaluated on three fruits per accession on a 1-5 scale by applying an ascending pressure with the extremity of the thumb on the fruit surface. Bruisedness, which represents the susceptibility of the fruit to mechanical damages, was evaluated by visual inspection of the fruits 4 h after harvest. Analyses were performed for two consecutive years except for FIRM and SR traits which were evaluated a single year in 2021. TA and TSS were evaluated from a homogenized pool of a minimum of 10 fruits with a pH-metric titration with sodium hydroxide of 10 g fruit puree and an Atago Handheld (PAL-1) Digital Pocket Refractometer (Atago, Saitama, Japan), respectively.

### Statistical analysis

Best Unbiased Linear Predictors (BLUPs) for the diversity panel were calculated using a linear mixed model (LMM) from the lme4 R package^68^:

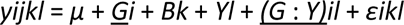

where Y/E represented the fixed effects of year/environments, B the fixed effect of blocks, G the random genotypic, GxY/E genotype x year/environment effects and e the residual effects.

Variance components for these effects were estimated using restricted maximum likelihood (REML). Broad sense heritability was estimated as follows:

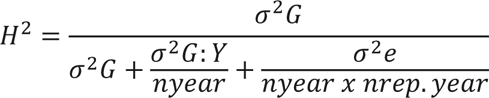

where genotype (G) variance at the numerator. Random variance components involving year (Y) were divided by the mean number of years (*nyear*). Other random variance components involving block effects or residuals were divided by the mean number of years times the mean number of replicates per year (*nrep*.*year*).

Pearson correlation between different traits were calculated using ‘cor’ function and visualized by ‘corrplot’ v. 0.92 R package. PCA on all traits was performed using the prcomp function from R core and visualized with fviz_pca function from factoextra v.1.0.7 package or ggplot2 package. The impact of the structure on each variable was assessed by simple regression of the genetic groups on their respective phenotypes.

### Genotyping

DNA was extracted from young leaves with a CTAB method adapted from Sánchez-Sevilla *et al.*, (2015)^33^. Samples were genotyped using Affymetrix® 50K FanaSNP array^21^ in the ‘Gentyane’ genotyping platform (Clermont-Auvergne-Rhône-Alpes, INRAE, France). SNP calling was processed through Axiom™ Analysis Suite software (v5.1.1.1; Thermo Fisher Scientific, Inc.) following the best practices of the software documentation. Accessions with missing data higher than 3% were removed from analysis. Markers presenting more than 5% of missing data and minor allele frequencies of less than 5% were filtered out.

### Structure and genetic diversity analysis

We performed a structure population analysis using STRUCTURE (v2.3.4^69^) with 5 runs for a range of K = 2 to 10 with 38,120 markers. The burn-in period length was set to 10,000 and 20,000 Markov Chain Monte Carlo (MCMC). The best fitting K was identified with STRUCTURE HARVESTER^70^. Plots were performed using the ggplot2 v.3.3.6 package^71^. PCA analyses were performed with PCA function from factorMinerR v.2.7 package^72^. Additionally, we included genotypes from Hardigan et al., (2021b)^4^ and Zurn et al. (2022)^27^ to perform PCA using the prcomp function from R core and visualize with fviz_pca_ind function from factoextra v.1.0.7 package or ggplot2 package. We conducted a ML tree with the 233 accessions using IQ-TREE v.2.1.3^73^ with 1000 bootstrap and the TVMe+ASC+R3 model suitable for SNP arrays. Linkage disequilibrium for each chromosome and genetic group was computed using the LDcorSV v.1.3.3 package^74^. Nucleotide diversity among each genetic group was calculated using TASSEL^75^. Finally, we performed principal component analysis-based genomes scans to detect markers under selection using the pcadapt package^76^, implementing the pcadapt function with K = 3. Output from genome scans were then compared with nucleotide diversity profiles to search for selective sweeps.

### Genome-Wide Association Study

The association mapping was performed using GAPIT v.3^77^ using the Camarosa genome physical positions^1^ with the Bayesian-information and linkage-disequilibrium iteratively nested keyway (BLINK) model^78^. In order to control for confounding effects, the structure was implemented for each trait in two different ways by 1) adding the previously calculated structure parameters as covariates or 2) fitting directly principal components from the principal component analysis using the PCA.total argument. Best models were selected based on genomic inflation factors, λ. The kinship was determined from the SNP data using the VanRaden mean algorithm. The analysis was performed on yearly and across two years Best Linear Unbiased Predictors (BLUPs), using a 5% Bonferroni threshold. Manhattan and Quantile-Quantile plots were plotted using the CMplot R package. Allelic effects for each significant marker were plotted on adjusted means using the ggplot2 R package.

## Author’s contributions

BD and AlP conceived and designed the experiments. AlP conducted hands-on experiments and data collection. AuP, JoP and JuP contributed to data collection. AlP, PRS, BD and CR analyzed the data. AlP, BD and CR wrote the original draft. All authors read and approved the final manuscript.

## Acknowledgements

We thank Sarah Touzani and Eva Bouillon for their help in phenotyping. The project was funded by the Nouvelle-Aquitaine Region and the European Regional Development Fund (ERDF) (REGINA project no. 67822110; AgirClim project No. 2018-1R20202); and European Union Horizon 2020 research and innovation program (BreedingValue project No. 101000747; PRIMA-Partnership 2019-2022 Med-Berry project). AlP was supported by the CIFRE (Convention Industrielle de Formation par la Recherche) contract between Invenio (SME, Bordeaux, France) and the INRAE BAP department.

## Data availability

All relevant data generated or analyzed are included in the manuscript and the supporting materials.

## Supplementary data

**Supplementary Figure S1.** Distribution of the year of release for the 223 accessions of the diversity panel.

**Supplementary Figure S2.** Phylogenetic tree of the 223 accessions of the diversity panel using the TVMe+ASC+R3 model.

**Supplementary Figure S3.** π chromosome-wide estimates for each genetic group for 400kb windows across the octoploid genome.

**Supplementary Figure S4.** Linkage disequilibrium (LD) decay along each chromosome of the octoploid genome.

**Supplementary Figure S5.** Distribution of the Invenio panel (green dots) among published data.

**Supplementary Figure S6.** Distribution of BLUP estimates for the 12 traits.

**Supplementary Figure S7**. Principal Component Analysis of the 2-year BLUP values for 11 traits.

**Supplementary Figure S8**. Correlations between the 12 traits for each year for each genetic group.

**Supplementary Figure S9**. Comparisons of 2-year BLUP values for FIRM, BRU, UFS, UCOL, COL, TA, TSS and BA among genetic groups.

**Supplementary Figure S10**. Genetic gains for FIRM, BRU, UFS, UCOL, TA, TSS and BA among genetic groups.

**Supplementary Figure S11.** Genome wide association of study of UFS and ACH.

**Supplementary Table S1.** List of the 223 genotypes.

**Supplementary Table S2.** Pearson correlations between 2-year estimated BLUP values for the 12 traits.

**Supplementary Table S3.** List of significant trait associations obtained for the GWAS on the 12 traits.

**Supplementary Table S4.** Genome scan outputs for the 71 trait associations.

**Supplementary Table S5.** Functions and/or possible roles of the 64 candidate genes underlying the trait associations.

