## Supplemental Figures S1-to-S11 and Tables S1-S2 for "Exploration of a European-centered strawberry diversity panel provides markers and candidate genes for the control of fruit quality traits"

Supplementary File

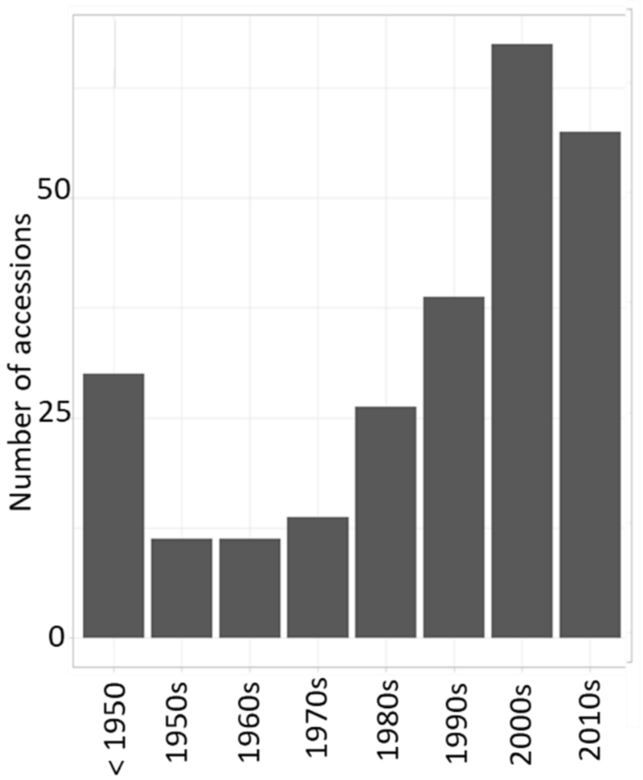


S**upplementary Figure S1**. Distribution of the year of release for the 223 accessions of the diversity panel.

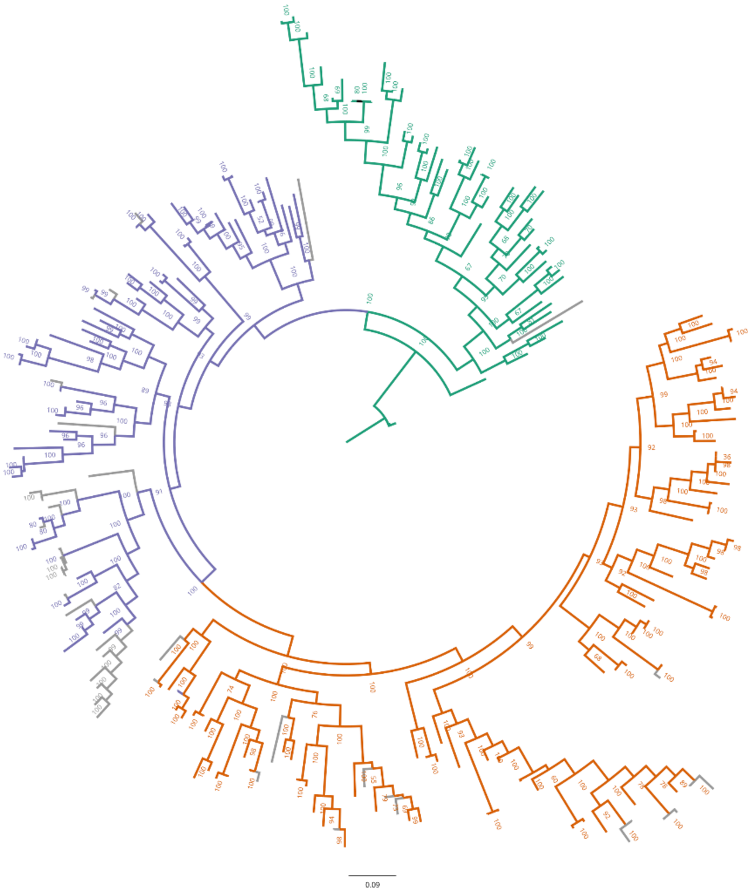


**Supplementary Figure S2**. Phylogenetic tree of the 223 accessions of the diversity panel using the TVMe+ASC+R3 model. The genetic group 1 is colored in green, group 2 in purple and group 3 in orange. Individuals clustered within the wrong group are labeled in grey. Groups 1, 2, 3: Heirloom & related, European mixed group and American & European mixed groups, respectively.

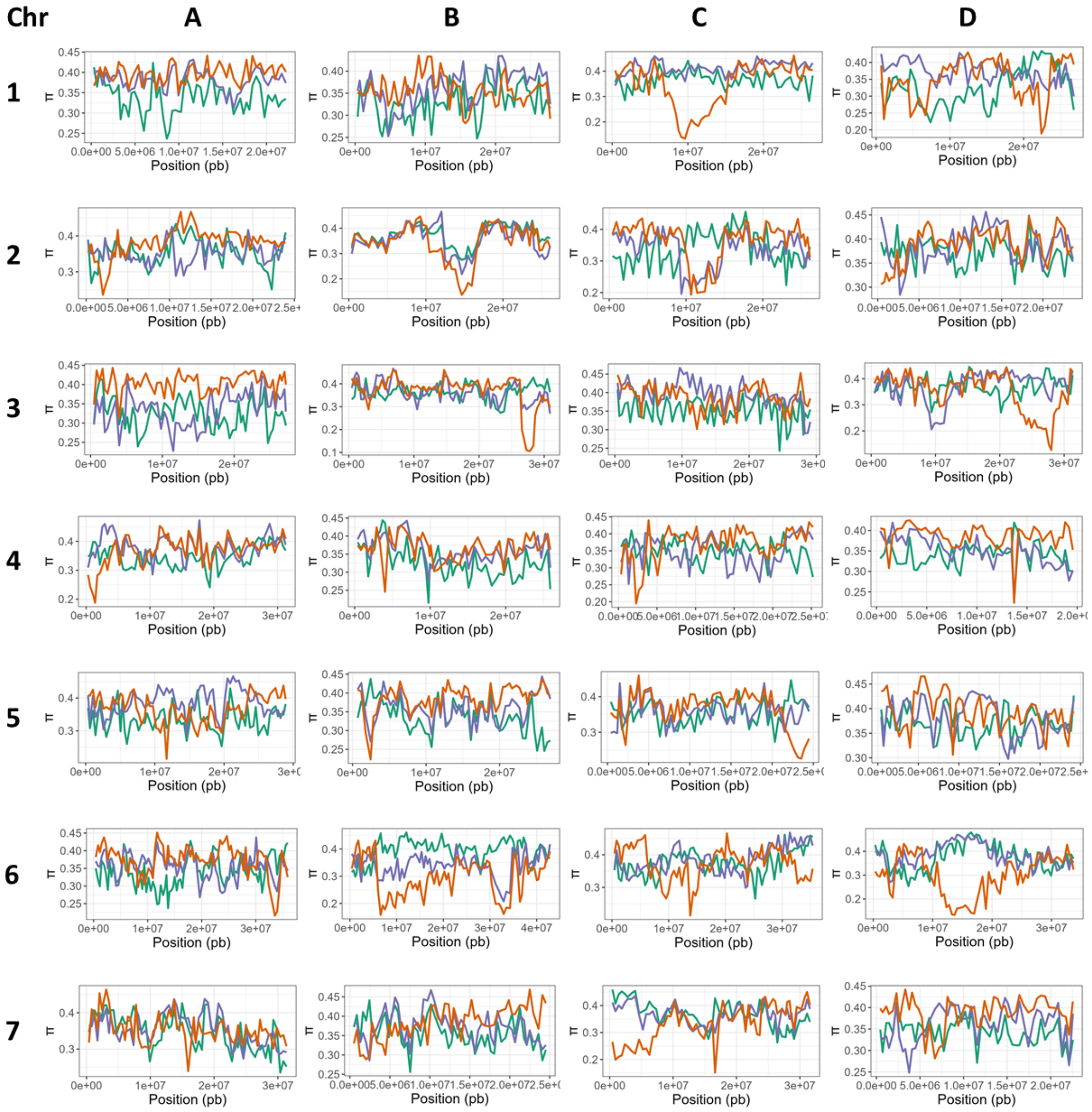


**Supplementary Figure S3**. π chromosome-wide estimates for each genetic group for 400kb windows across the octoploid genome. The genetic group 1 is colored in green, group 2 in purple and group 3 in orange. Groups 1, 2, 3: Heirloom & related, European mixed group and American & European mixed groups, respectively.

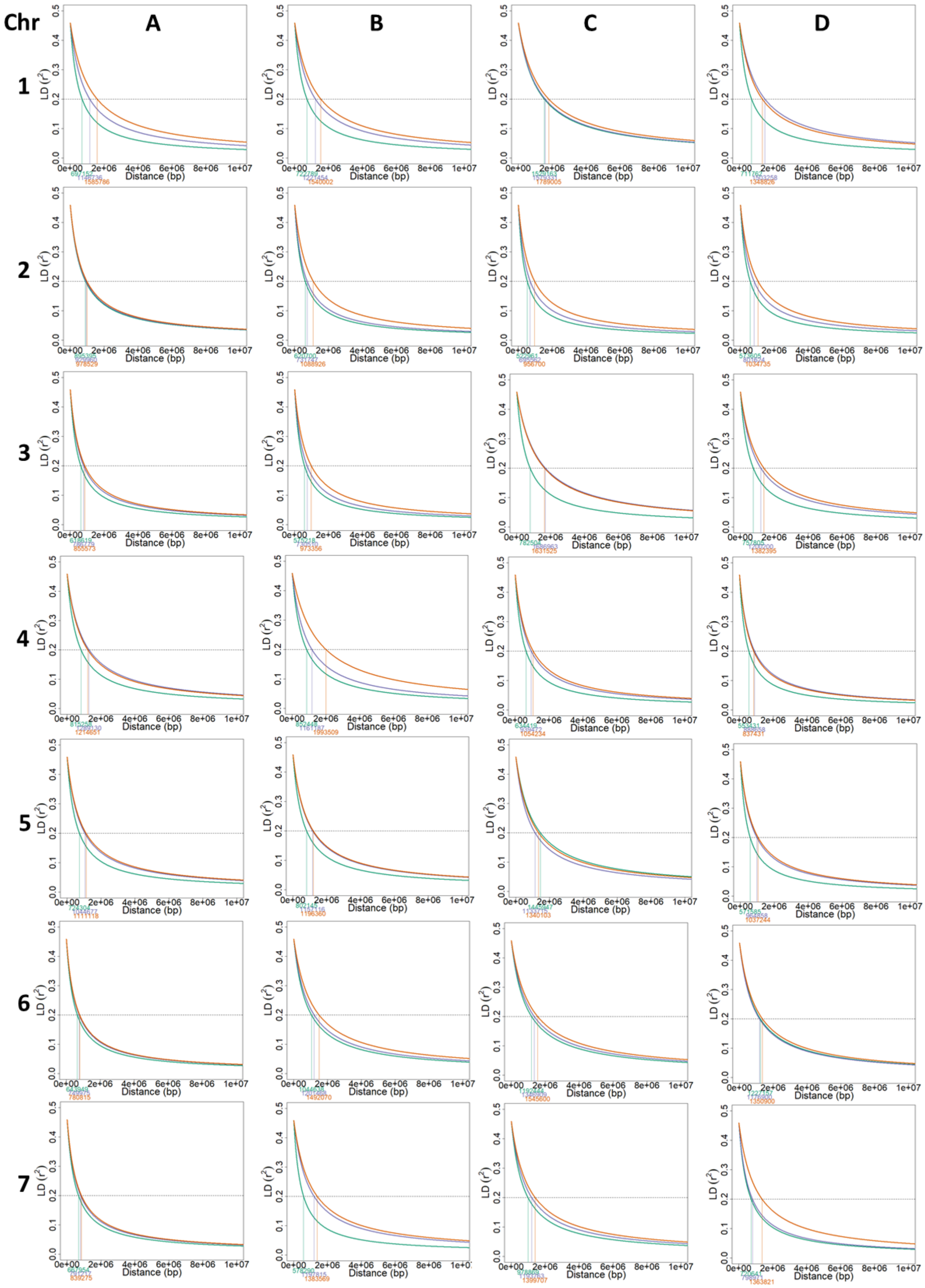
Chr
A
B
C
D
1
2
3
4
5
6
7
LD (r²)
Distance (bp)

**Supplementary Figure S4**. Linkage disequilibrium (LD) decay along each chromosome of the octoploid genome. The genetic group 1 is colored in green, group 2 in purple and group 3 in orange. The dashed line represents the LD decay at r2 = 0.2. Groups 1, 2, 3: Heirloom & related, European mixed group and American & European mixed groups, respectively

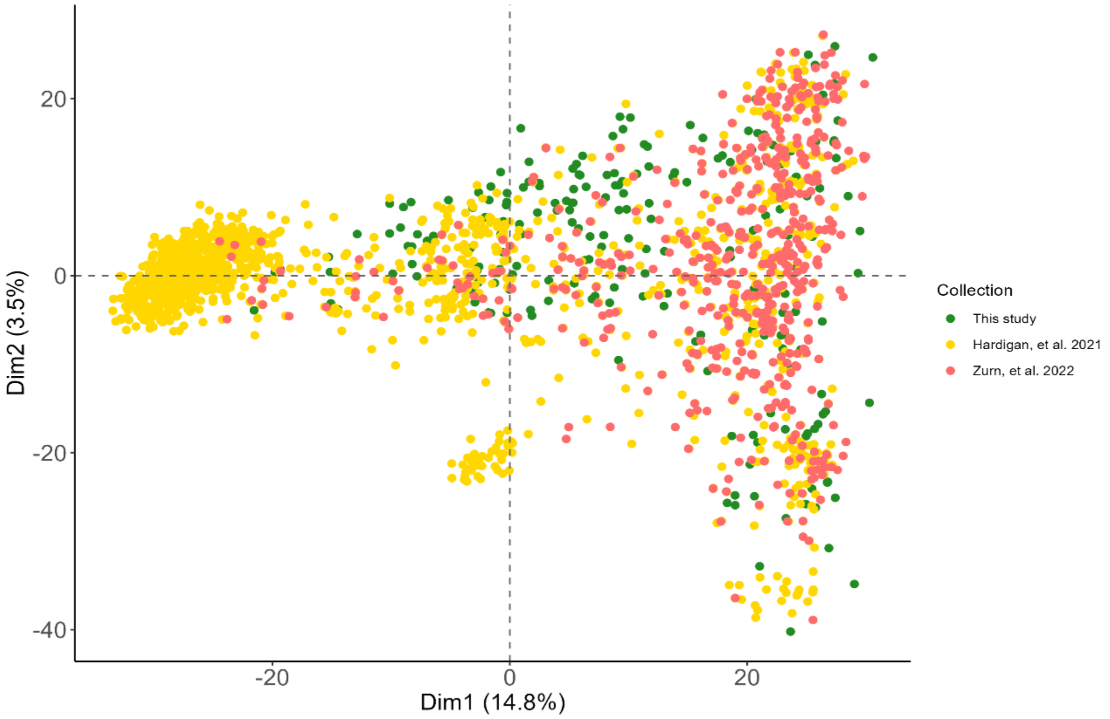


**Supplementary Figure S5**. Distribution of the Invenio panel (green dots) among published data. 1 569 genotypes (yellow dots) studied in Hardigan et al. (2020) and 539 genotypes studied in Zurn et al. (2022) (red dots) with 3 215 SNP markers.

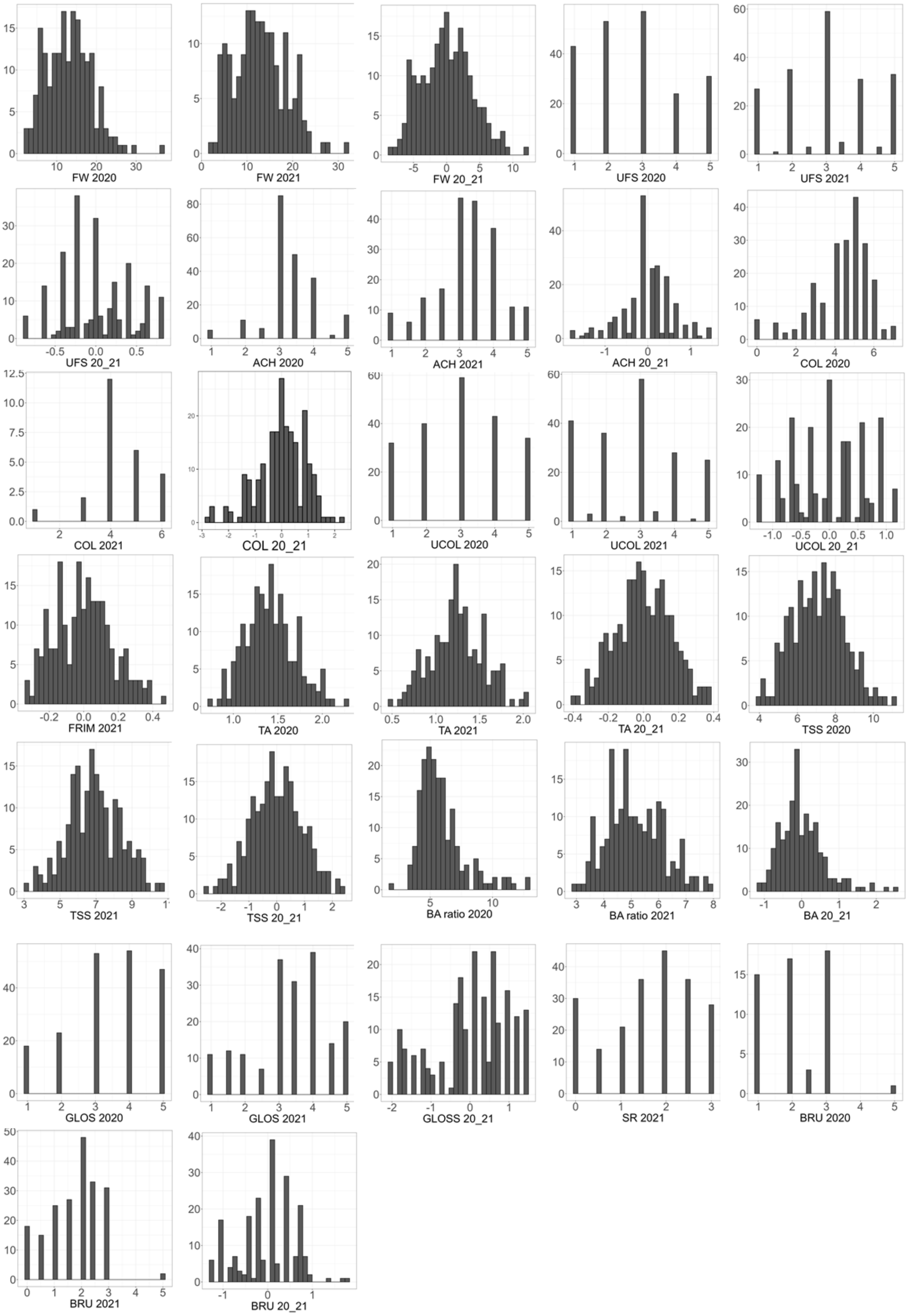
FW 2020
FW 2021
FW 20_21
UFS 2020
UFS 2021
UFS 20_21
ACH 2020
ACH 2021
ACH 20_21
COL 2020
COL 2021
COL 20_21
UCOL 2020
UCOL 2021
UCOL 20_21
FRIM 2021
TA 2020
TA 2021
TA 20_21
TSS 2020
TSS 2021
TSS 20_21
BA ratio 2020
BA ratio 2021
BA 20_21
GLOS 2020
GLOS 2021
GLOSS 20_21
SR 2021
BRU 2020
BRU 2021
BRU 20_21

**Supplementary Figure S6**. Distribution of BLUP estimates for the 12 traits. FW, fruit weight; UFS, uniformity of fruit shape; COL, skin color; UCOL, uniformity of skin color; ACH, position and depth of achenes; FIRM, firmness; TA, titratable acidity; TSS, total soluble solids; BA, Brix/TA ratio; GLOS, glossiness; SR, skin resistance; BRU, bruisedness.

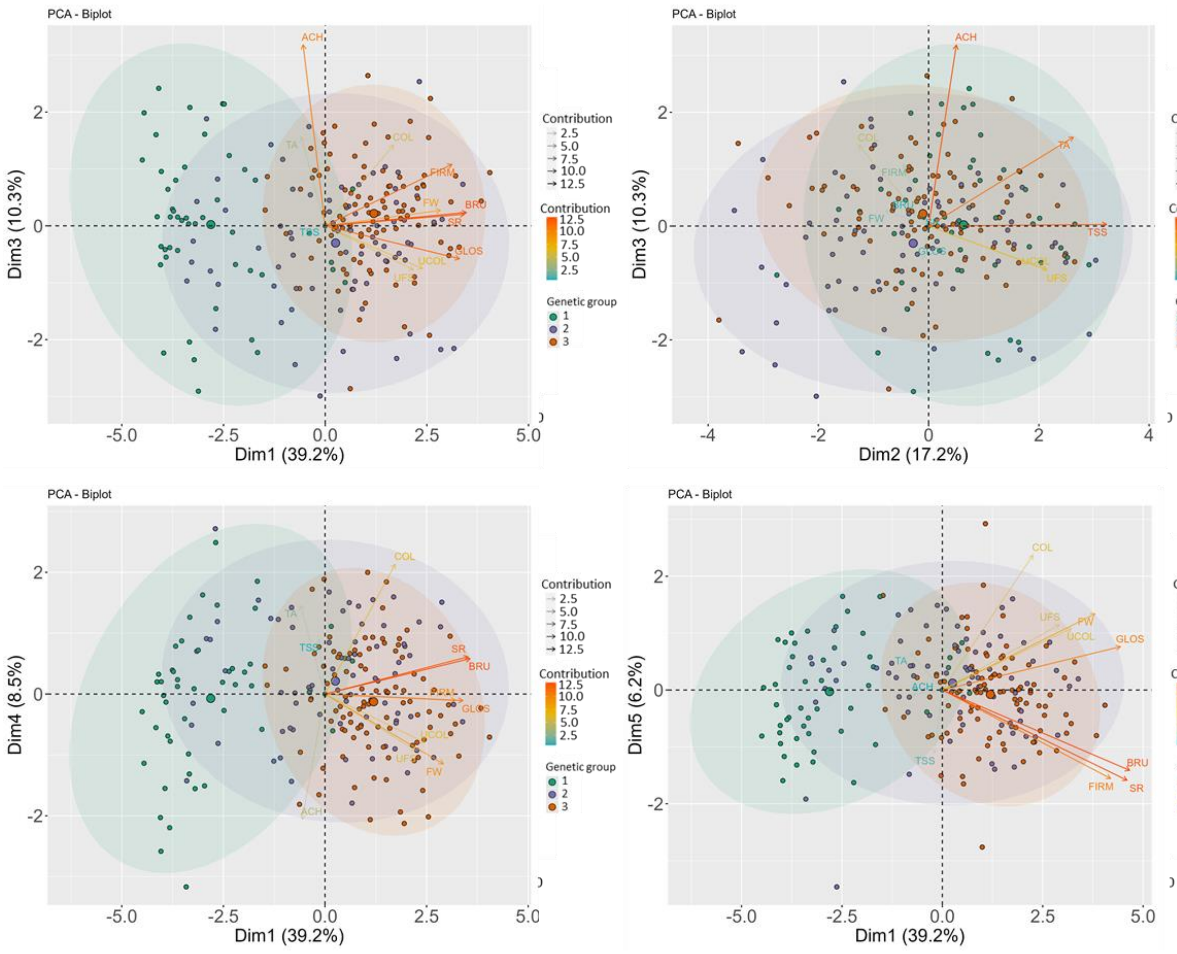

**Supplementary Figure S7**. Principal Component Analysis of the 2-year BLUP values for 11 traits. FW, fruit weight; UFS, uniformity of fruit shape; COL, skin color; UCOL, uniformity of skin color; ACH, position and depth of achenes; FIRM, firmness; TA, titratable acidity; TSS, total soluble solids; GLOS, glossiness; SR, skin resistance; BRU, bruisedness.

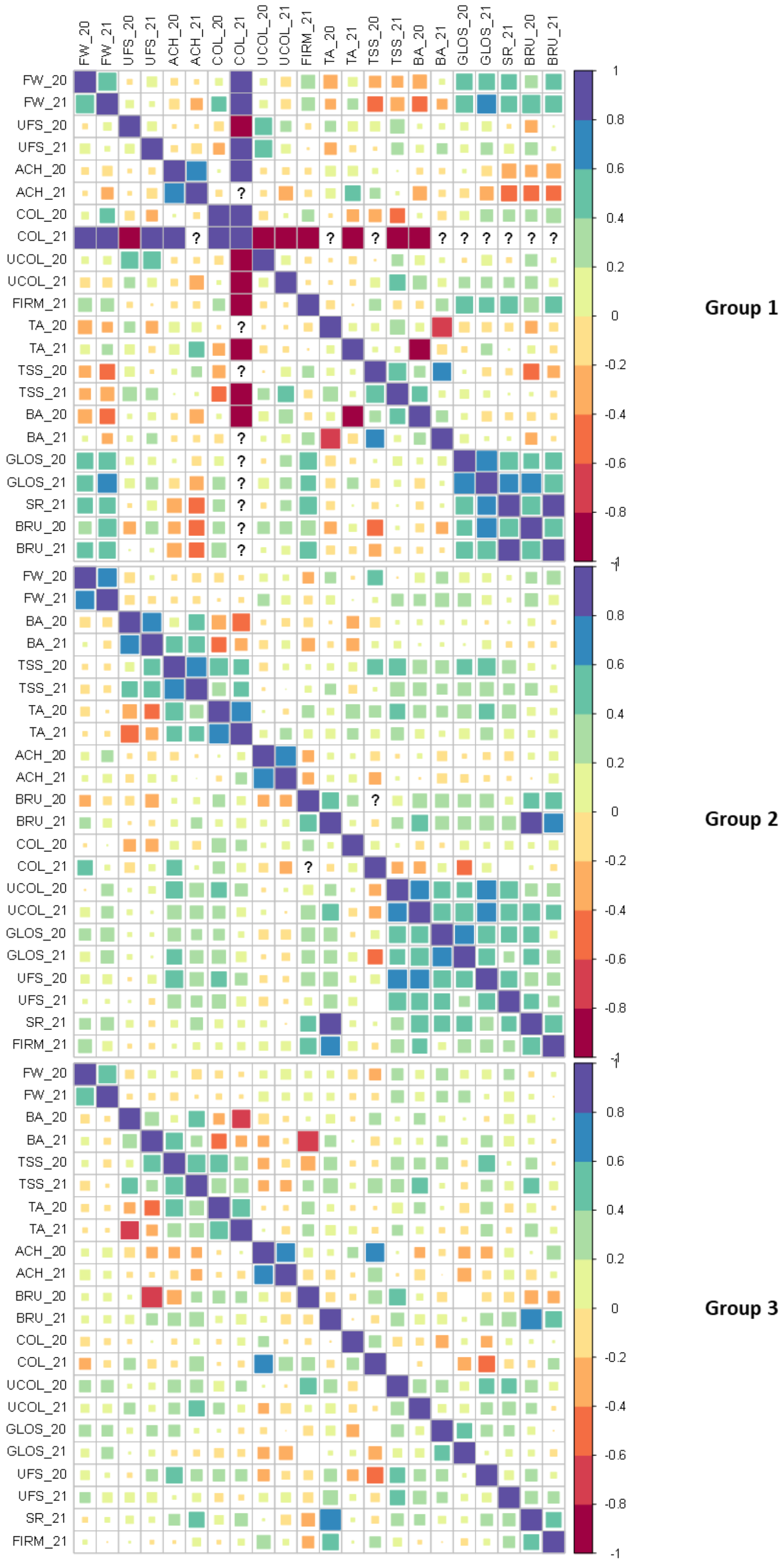


**Supplementary Figure S8**. Correlations between the 12 traits for each year for each genetic group. '?' when correlation could not be calculated.

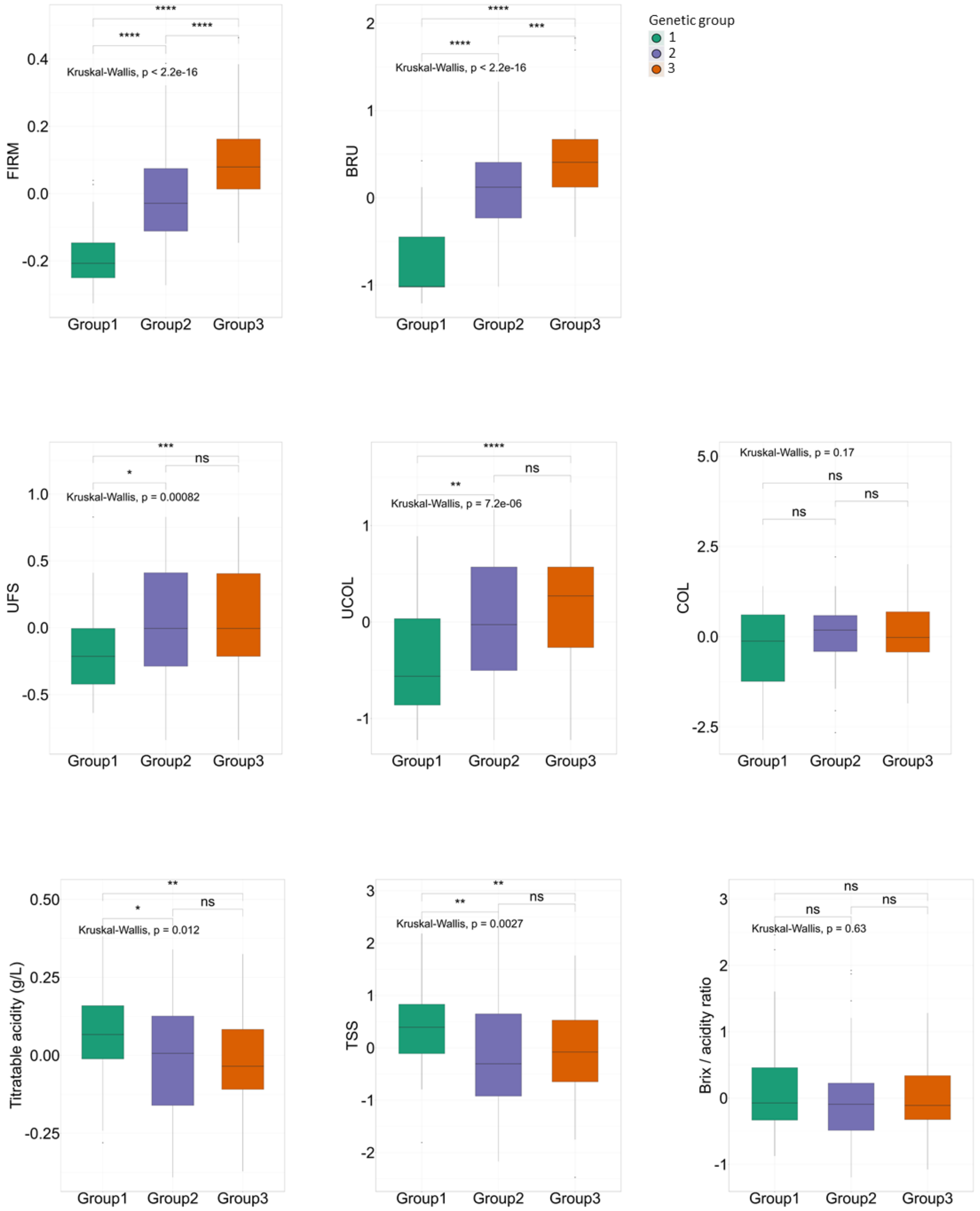


**Supplementary Figure S9**. Comparisons of 2-year BLUP values for FIRM, BRU, UFS, UCOL, COL, TA, TSS and BA among genetic groups. Genetic groups 1, 2 and 3 are colored in green, purple and orange, respectively. UFS, uniformity of fruit shape; COL, skin color; UCOL, uniformity of skin color; FIRM, firmness; TA, titratable acidity; TSS, total soluble solids; BRU, bruisedness.

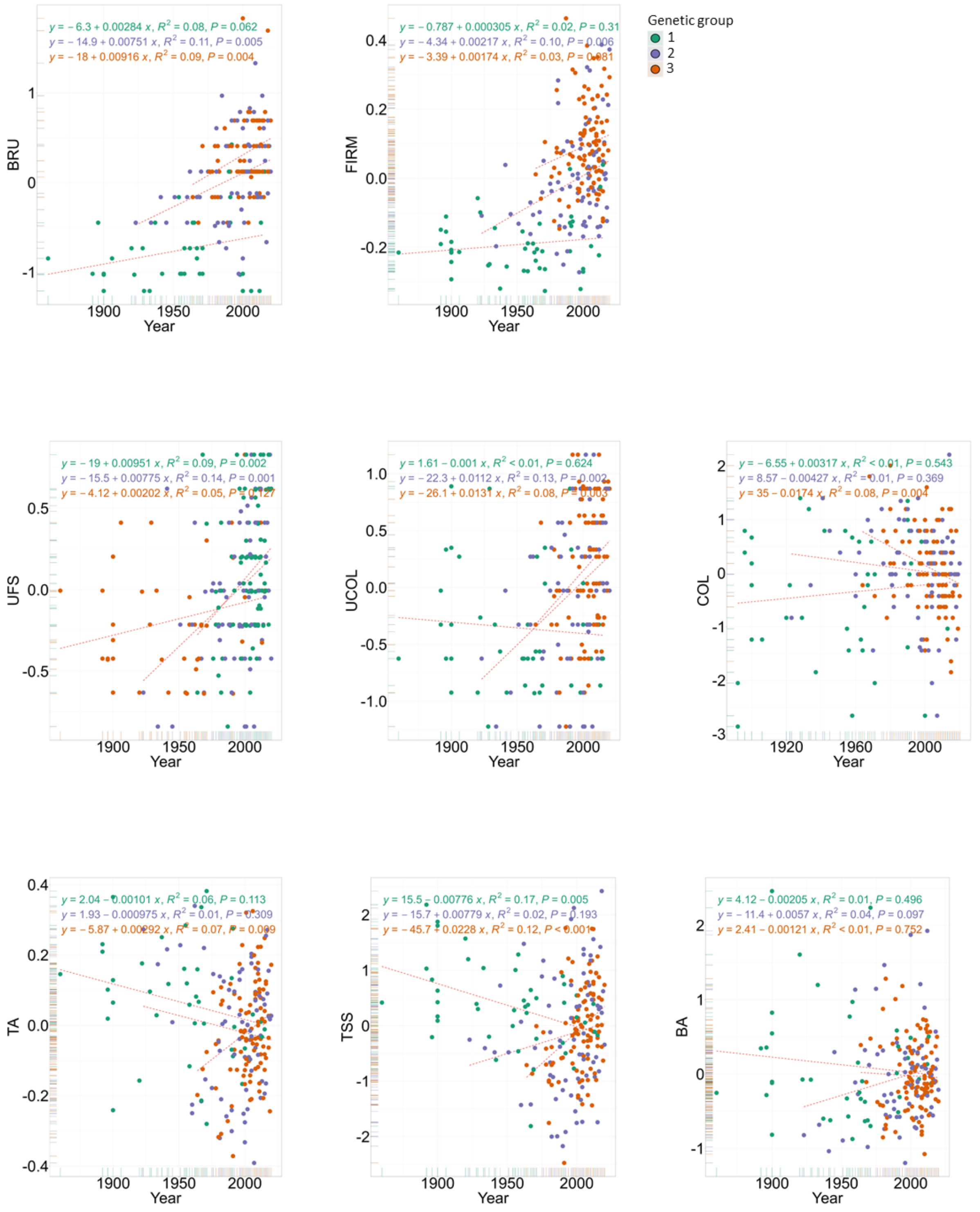


**Supplementary Figure S10**. Genetic gains for FIRM, BRU, UFS, UCOL, TA, TSS and BA among genetic groups. Genetic groups 1, 2 and 3 are colored in green, purple and orange, respectively. UFS, uniformity of fruit shape; COL, skin color; UCOL, uniformity of skin color; FIRM, firmness; TA, titratable acidity; TSS, total soluble solids; BRU, bruisedness.

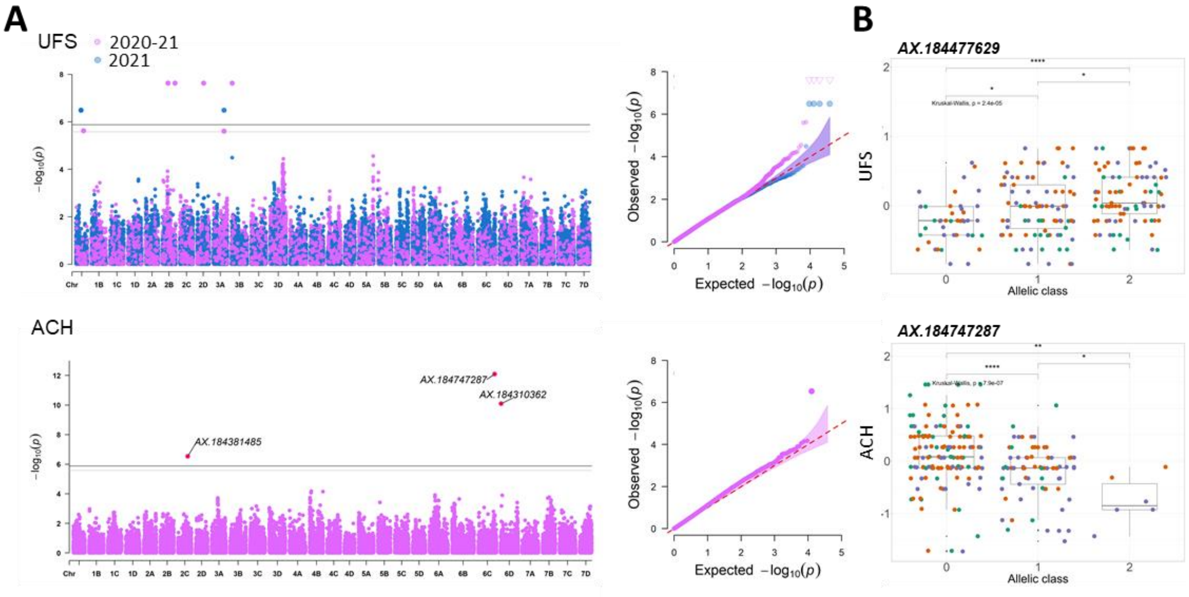


**Supplementary Figure S11**. Genome wide association of study of UFS and ACH. (A) Manhattan and Q-Q plots for yearly and 2-year BLUP values. (B) Effect of the most significant SNP markers. Genetic groups 1, 2 and 3 are colored in green, purple and orange, respectively. UFS, uniformity of fruit shape; ACH, position and depth of achenes. Marker classes are as follows: 0=AA genotype, 1=AB, and 2=BB genotype according to the Axiom™ Strawberry FanaSNP 50k.

**Supplementary Table S1**. List of the 223 genotypes. Origin (country/state), Continent, Year of release, group of structure and their estimated membership fractions (%). Groups were identified using Structure. Origins and dates of release/observation from Cost836, GenBerry database, CPOV, UC Davis.

| Name | Origin1 | Origin2 | Year | Group | Estimated membership fractions | | |
|---|---|---|---|---|---|---|---|
| | | | | | Group 1 | Group 2 | Group 3 |
| RG001 | Italy | Europe | 1982 | European mixed | 0.26 | 0.61 | 0.12 |
| RG002 | France | Europe | 1992 | Heirloom & related | 0.36 | 0.34 | 0.30 |
| RG004 | Japan | Asia | 1992 | Heirloom & related | 1.00 | 0.00 | 0.00 |
| RG005 | Italy | Europe | 2002 | American & European mixed | 0.00 | 0.45 | 0.55 |
| RG006 | Univ.Calif. | USA | 2004 | American & European mixed | 0.00 | 0.00 | 1.00 |
| RG008 | Maryland | America | 1981 | European mixed | 0.09 | 0.86 | 0.05 |
| RG009 | NA | NA | 2019 | American & European mixed | 0.15 | 0.20 | 0.65 |
| RG010 | France | Europe | 2012 | American & European mixed | 0.19 | 0.00 | 0.81 |
| RG011 | Italy | Europe | 2020 | European mixed | 0.02 | 0.59 | 0.39 |
| RG012 | Italy | Europe | 2020 | American & European mixed | 0.00 | 0.33 | 0.67 |
| RG013 | France | Europe | 2008 | European mixed | 0.24 | 0.53 | 0.23 |
| RG015 | France | Europe | 1998 | American & European mixed | 0.04 | 0.39 | 0.57 |
| RG016 | France | Europe | 2003 | American & European mixed | 0.24 | 0.15 | 0.61 |
| RG017 | France | Europe | 2012 | Heirloom & related | 1.00 | 0.00 | 0.00 |
| RG018 | Italy | Europe | 2018 | American & European mixed | 0.00 | 0.49 | 0.51 |
| RG019 | Maryland | America | 1981 | European mixed | 0.16 | 0.77 | 0.07 |
| RG020 | Univ.Calif. | America | 1997 | American & European mixed | 0.00 | 0.00 | 1.00 |
| RG021 | Italy | Europe | 2005 | American & European mixed | 0.02 | 0.43 | 0.55 |
| RG022 | UK | Europe | 2001 | American & European mixed | 0.31 | 0.12 | 0.57 |
| RG023 | Univ.Calif. | America | 1998 | Heirloom & related | 0.86 | 0.08 | 0.06 |
| RG024 | Belgium | Europe | 2000 | European mixed | 0.47 | 0.53 | 0.00 |
| RG026 | France | Europe | 1958 | Heirloom & related | 0.83 | 0.17 | 0.00 |
| RG027 | France | Europe | 1962 | European mixed | 0.04 | 0.96 | 0.00 |
| RG028 | France | Europe | 2008 | American & European mixed | 0.09 | 0.13 | 0.78 |
| RG029 | Maryland | America | 1923 | European mixed | 0.13 | 0.87 | 0.00 |
| RG030 | Germany | Europe | 1900 | Heirloom & related | 1.00 | 0.00 | 0.00 |
| RG031 | France | Europe | 2000 | Heirloom & related | 1.00 | 0.00 | 0.00 |
| RG032 | Netherlands | Europe | 1971 | American & European mixed | 0.31 | 0.04 | 0.65 |
| RG033 | Univ.Calif. | America | 1978 | American & European mixed | 0.15 | 0.10 | 0.75 |
| RG034 | France | Europe | 2018 | American & European mixed | 0.00 | 0.14 | 0.86 |
| RG035 | Univ Calif | USA | 1992 | American & European mixed | 0.00 | 0.00 | 1.00 |
| RG036 | France | Europe | 2008 | European mixed | 0.03 | 0.73 | 0.23 |
| RG037 | Spain | Europe | 2003 | American & European mixed | 0.00 | 0.00 | 1.00 |
| RG038 | France | Europe | 2001 | American & European mixed | 0.06 | 0.15 | 0.79 |
| RG039 | Univ.Calif. | America | 1989 | American & European mixed | 0.10 | 0.03 | 0.86 |
| RG041 | France | Europe | 2010 | American & European mixed | 0.00 | 0.42 | 0.58 |
| RG043 | New-York | America | 1934 | European mixed | 0.34 | 0.54 | 0.12 |
| RG044 | France | Europe | 1989 | American & European mixed | 0.00 | 0.43 | 0.57 |
| RG045 | France | Europe | 1991 | Heirloom & related | 0.69 | 0.16 | 0.16 |
| RG047 | France | Europe | 1992 | American & European mixed | 0.00 | 0.31 | 0.69 |
| RG048 | France | Europe | 2018 | European mixed | 0.17 | 0.48 | 0.34 |
| RG049 | France | Europe | 2019 | European mixed | 0.02 | 0.63 | 0.34 |
| RG050 | France | Europe | 2018 | European mixed | 0.00 | 0.65 | 0.35 |
| RG052 | France | Europe | 2004 | European mixed | 0.06 | 0.63 | 0.31 |
| RG053 | France | Europe | 2004 | American & European mixed | 0.01 | 0.47 | 0.52 |
| RG055 | France | Europe | 2005 | European mixed | 0.00 | 0.63 | 0.37 |
| RG056 | France | Europe | 2005 | European mixed | 0.14 | 0.84 | 0.01 |
| RG057 | France | Europe | 2005 | European mixed | 0.02 | 0.93 | 0.05 |
| RG058 | France | Europe | 2005 | American & European mixed | 0.37 | 0.18 | 0.45 |
| RG059 | France | Europe | 2005 | American & European mixed | 0.23 | 0.06 | 0.71 |
| RG060 | France | Europe | 2007 | European mixed | 0.34 | 0.41 | 0.25 |
| RG063 | France | Europe | 2008 | American & European mixed | 0.00 | 0.47 | 0.52 |
| RG065 | France | Europe | 2008 | American & European mixed | 0.03 | 0.39 | 0.58 |
| RG066 | France | Europe | 2009 | European mixed | 0.00 | 0.58 | 0.42 |
| RG067 | France | Europe | 2010 | European mixed | 0.00 | 0.82 | 0.18 |
| RG068 | France | Europe | 2010 | American & European mixed | 0.23 | 0.01 | 0.76 |
| RG069 | France | Europe | 2010 | American & European mixed | 0.00 | 0.37 | 0.63 |
| RG071 | France | Europe | 2010 | American & European mixed | 0.04 | 0.35 | 0.61 |
| RG072 | France | Europe | 2010 | American & European mixed | 0.12 | 0.25 | 0.62 |
| RG073 | France | Europe | 2011 | American & European mixed | 0.00 | 0.35 | 0.65 |
| RG074 | France | Europe | 2011 | American & European mixed | 0.05 | 0.28 | 0.67 |
| RG075 | France | Europe | 2012 | American & European mixed | 0.05 | 0.24 | 0.70 |
| RG076 | France | Europe | 2012 | European mixed | 0.00 | 0.59 | 0.40 |
| RG077 | France | Europe | 2015 | American & European mixed | 0.00 | 0.45 | 0.55 |
| RG078 | France | Europe | 2015 | American & European mixed | 0.08 | 0.38 | 0.54 |
| RG079 | France | Europe | 2015 | American & European mixed | 0.12 | 0.35 | 0.53 |

### Table S1 - to be continued

| | | | | | | | |
|---|---|---|---|---|---|---|---|
| RG081 | Univ.Calif. | America | 1983 | American & European mixed | 0.00 | 0.00 | 1.00 |
| RG082 | France | Europe | 2000 | American & European mixed | 0.12 | 0.16 | 0.71 |
| RG084 | Chile | America | NA | Heirloom & related | 1.00 | 0.00 | 0.00 |
| RG085 | Chile | America | NA | Heirloom & related | 0.79 | 0.21 | 0.00 |
| RG086 | Netherlands | Europe | 2002 | European mixed | 0.04 | 0.65 | 0.30 |
| RG087 | France | Europe | 1998 | European mixed | 0.03 | 0.79 | 0.18 |
| RG088 | France | Europe | 1996 | American & European mixed | 0.00 | 0.48 | 0.52 |
| RG089 | France | Europe | 1999 | American & European mixed | 0.14 | 0.32 | 0.54 |
| RG090 | France | Europe | 1996 | European mixed | 0.02 | 0.98 | 0.00 |
| RG091 | France | Europe | 1996 | European mixed | 0.01 | 0.63 | 0.37 |
| RG092 | France | Europe | 2001 | American & European mixed | 0.25 | 0.04 | 0.72 |
| RG093 | France | Europe | 1998 | European mixed | 0.05 | 0.95 | 0.00 |
| RG094 | France | Europe | 2010 | American & European mixed | 0.20 | 0.07 | 0.72 |
| RG095 | France | Europe | 2010 | American & European mixed | 0.00 | 0.05 | 0.95 |
| RG096 | France | Europe | 2011 | American & European mixed | 0.00 | 0.34 | 0.66 |
| RG097 | France | Europe | 2011 | American & European mixed | 0.21 | 0.14 | 0.65 |
| RG098 | France | Europe | 2012 | American & European mixed | 0.00 | 0.47 | 0.53 |
| RG099 | France | Europe | 2012 | American & European mixed | 0.23 | 0.08 | 0.69 |
| RG100 | France | Europe | 2014 | American & European mixed | 0.00 | 0.26 | 0.74 |
| RG101 | France | Europe | 2014 | American & European mixed | 0.22 | 0.01 | 0.78 |
| RG102 | France | Europe | 2015 | American & European mixed | 0.02 | 0.29 | 0.70 |
| RG104 | France | Europe | 1997 | American & European mixed | 0.25 | 0.02 | 0.73 |
| RG105 | France | Europe | 1997 | American & European mixed | 0.28 | 0.15 | 0.57 |
| RG106 | France | Europe | 1996 | American & European mixed | 0.00 | 0.44 | 0.56 |
| RG107 | Italy | Europe | 2002 | European mixed | 0.00 | 0.63 | 0.36 |
| RG108 | Italy | Europe | 2013 | European mixed | 0.11 | 0.88 | 0.02 |
| RG109 | France | Europe | 2004 | European mixed | 0.06 | 0.71 | 0.23 |
| RG110 | France | Europe | 1996 | European mixed | 0.00 | 0.54 | 0.46 |
| RG111 | Florida | America | 1992 | American & European mixed | 0.00 | 0.33 | 0.67 |
| RG112 | France | Europe | 2012 | American & European mixed | 0.00 | 0.43 | 0.57 |
| RG113 | France | Europe | 2011 | European mixed | 0.00 | 0.67 | 0.33 |
| RG114 | France | Europe | 2016 | American & European mixed | 0.00 | 0.38 | 0.62 |
| RG115 | Italy | Europe | 2014 | European mixed | 0.03 | 0.53 | 0.44 |
| RG117 | France | Europe | 2007 | American & European mixed | 0.00 | 0.46 | 0.54 |
| RG119 | France | Europe | 2006 | American & European mixed | 0.02 | 0.21 | 0.77 |
| RG120 | Univ.Calif. | America | 1945 | European mixed | 0.14 | 0.60 | 0.26 |
| RG121 | Florida | America | 1980 | American & European mixed | 0.14 | 0.39 | 0.47 |
| RG122 | France | Europe | 2012 | American & European mixed | 0.04 | 0.22 | 0.74 |
| RG123 | Poland | Europe | 1985 | European mixed | 0.36 | 0.62 | 0.02 |
| RG124 | Maryland | America | 1975 | European mixed | 0.00 | 0.99 | 0.01 |
| RG125 | UK | Europe | 2008 | European mixed | 0.16 | 0.70 | 0.14 |
| RG127 | Netherlands | Europe | 1981 | European mixed | 0.00 | 1.00 | 0.00 |
| RG128 | Italy | Europe | 1970 | European mixed | 0.05 | 0.50 | 0.45 |
| RG129 | Netherlands | Europe | 1967 | European mixed | 0.39 | 0.61 | 0.00 |
| RG130 | France | Europe | 2000 | Heirloom & related | 1.00 | 0.00 | 0.00 |
| RG131 | UK | Europe | 2006 | American & European mixed | 0.12 | 0.10 | 0.78 |
| RG132 | France | Europe | 1976 | American & European mixed | 0.22 | 0.37 | 0.41 |
| RG133 | France | Europe | 1900 | Heirloom & related | 0.66 | 0.34 | 0.00 |
| RG134 | Florida | America | 2000 | American & European mixed | 0.00 | 0.04 | 0.96 |
| RG135 | Italy | Europe | 2018 | American & European mixed | 0.00 | 0.04 | 0.96 |
| RG138 | UK | Europe | 1990 | Heirloom & related | 0.99 | 0.00 | 0.01 |
| RG139 | Norway | Europe | 2001 | European mixed | 0.09 | 0.65 | 0.26 |
| RG140 | Italy | Europe | 1991 | American & European mixed | 0.25 | 0.37 | 0.38 |
| RG141 | France | Europe | 1976 | European mixed | 0.13 | 0.85 | 0.01 |
| RG142 | Germany | Europe | 1967 | Heirloom & related | 0.56 | 0.33 | 0.11 |
| RG143 | Germany | Europe | 1941 | European mixed | 0.17 | 0.71 | 0.12 |
| RG145 | Switzerland | Europe | 1990 | European mixed | 0.15 | 0.85 | 0.00 |
| RG146 | France | Europe | 2015 | European mixed | 0.20 | 0.74 | 0.06 |
| RG147 | Netherlands | Europe | 1960 | European mixed | 0.06 | 0.94 | 0.00 |
| RG148 | Germany | Europe | 1967 | Heirloom & related | 0.64 | 0.29 | 0.06 |
| RG149 | NA | NA | 1900 | Heirloom & related | 0.91 | 0.01 | 0.09 |
| RG150 | France | Europe | 2010 | European mixed | 0.01 | 0.51 | 0.48 |
| RG151 | France | Europe | 2009 | American & European mixed | 0.12 | 0.43 | 0.46 |
| RG152 | France | Europe | 1928 | Heirloom & related | 0.90 | 0.10 | 0.00 |
| RG153 | Germany | Europe | 1971 | Heirloom & related | 0.84 | 0.16 | 0.00 |
| RG158 | New-York | America | 1985 | European mixed | 0.21 | 0.79 | 0.00 |
| RG159 | Netherlands | Europe | 2015 | Heirloom & related | 1.00 | 0.00 | 0.00 |
| RG160 | Ukraine | Europe | 1900 | Heirloom & related | 0.94 | 0.04 | 0.02 |
| RG162 | Italy | Europe | 2001 | American & European mixed | 0.00 | 0.04 | 0.96 |
| RG164 | Canada | America | 1962 | Heirloom & related | 0.81 | 0.15 | 0.03 |
| RG166 | Maryland | America | 1969 | European mixed | 0.37 | 0.50 | 0.13 |
| RG167 | Italy | Europe | 2011 | American & European mixed | 0.01 | 0.15 | 0.84 |
| RG168 | France | Europe | 1896 | Heirloom & related | 0.54 | 0.46 | 0.00 |
| RG169 | France | Europe | 1900 | Heirloom & related | 0.72 | 0.24 | 0.03 |

Table S1 - to be continued

| | | | | | | | |
|---|---|---|---|---|---|---|---|
| RG170 | France | Europe | 1906 | Heirloom & related | 0.91 | 0.01 | 0.08 |
| RG171 | France | Europe | 2016 | European mixed | 0.14 | 0.65 | 0.20 |
| RG174 | UK | Europe | 2012 | European mixed | 0.00 | 0.54 | 0.46 |
| RG175 | UK | Europe | 2013 | American & European mixed | 0.09 | 0.18 | 0.72 |
| RG176 | France | Europe | 2005 | European mixed | 0.37 | 0.38 | 0.25 |
| RG177 | France | Europe | 1991 | Heirloom & related | 0.48 | 0.23 | 0.30 |
| RG178 | France | Europe | 1955 | Heirloom & related | 0.90 | 0.05 | 0.04 |
| RG179 | France | Europe | 2016 | American & European mixed | 0.21 | 0.06 | 0.73 |
| RG180 | Belgium | Europe | 1986 | European mixed | 0.34 | 0.66 | 0.00 |
| RG181 | UK | Europe | 1965 | Heirloom & related | 0.44 | 0.13 | 0.43 |
| RG182 | Germany | Europe | 1933 | Heirloom & related | 0.95 | 0.03 | 0.01 |
| RG184 | Univ.Calif. | America | 2008 | American & European mixed | 0.02 | 0.00 | 0.98 |
| RG185 | Japan | Asia | 1960 | European mixed | 0.21 | 0.67 | 0.12 |
| RG186 | France | Europe | 1971 | Heirloom & related | 0.88 | 0.09 | 0.02 |
| RG187 | Italy | Europe | 2015 | American & European mixed | 0.00 | 0.47 | 0.53 |
| RG188 | Russia | Europe | 1958 | Heirloom & related | 0.76 | 0.16 | 0.08 |
| RG189 | Italy | Europe | 2000 | American & European mixed | 0.04 | 0.03 | 0.93 |
| RG190 | Italy | Europe | 2001 | American & European mixed | 0.01 | 0.12 | 0.87 |
| RG191 | Japan | Asia | 1984 | European mixed | 0.32 | 0.43 | 0.25 |
| RG192 | France | Europe | 2017 | European mixed | 0.02 | 0.50 | 0.48 |
| RG193 | Univ.Calif. | America | 1987 | American & European mixed | 0.00 | 0.00 | 1.00 |
| RG194 | Netherlands | Europe | 1969 | Heirloom & related | 0.64 | 0.27 | 0.09 |
| RG195 | Univ.Calif. | America | 1978 | American & European mixed | 0.01 | 0.17 | 0.82 |
| RG196 | Italy | Europe | 1997 | American & European mixed | 0.00 | 0.17 | 0.83 |
| RG198 | Italy | Europe | 1999 | European mixed | 0.08 | 0.84 | 0.08 |
| RG199 | UK | Europe | 1990 | European mixed | 0.29 | 0.71 | 0.00 |
| RG201 | Netherlands | Europe | 1980 | American & European mixed | 0.00 | 0.00 | 1.00 |
| RG202 | Univ.Calif. | America | 2008 | American & European mixed | 0.06 | 0.00 | 0.94 |
| RG204 | Denmark | Europe | 1990 | Heirloom & related | 0.63 | 0.31 | 0.06 |
| RG205 | NA | NA | 1990 | American & European mixed | 0.00 | 0.30 | 0.70 |
| RG207 | Italy | Europe | 2003 | European mixed | 0.10 | 0.86 | 0.04 |
| RG208 | Netherlands | Europe | 1964 | Heirloom & related | 0.72 | 0.26 | 0.02 |
| RG209 | Japan | Asia | 1990 | Heirloom & related | 0.45 | 0.32 | 0.23 |
| RG210 | Germany | Europe | 2017 | European mixed | 0.27 | 0.54 | 0.19 |
| RG211 | Netherlands | Europe | 1956 | Heirloom & related | 0.89 | 0.11 | 0.00 |
| RG212 | Germany | Europe | 1920 | Heirloom & related | 0.43 | 0.37 | 0.20 |
| RG214 | Florida | America | 1999 | American & European mixed | 0.03 | 0.21 | 0.76 |
| RG216 | UK | Europe | 1892 | Heirloom & related | 0.94 | 0.00 | 0.06 |
| RG217 | France | Europe | 2016 | European mixed | 0.00 | 0.51 | 0.49 |
| RG218 | Australia | Australia | 2004 | American & European mixed | 0.00 | 0.08 | 0.92 |
| RG222 | UK | Europe | 2008 | American & European mixed | 0.00 | 0.42 | 0.58 |
| RG223 | France | Europe | 1892 | Heirloom & related | 0.62 | 0.34 | 0.04 |
| RG224 | Canada | America | 2004 | American & European mixed | 0.07 | 0.39 | 0.53 |
| RG225 | Univ.Calif. | America | 2008 | American & European mixed | 0.02 | 0.00 | 0.98 |
| RG227 | France | Europe | 1937 | Heirloom & related | 0.99 | 0.00 | 0.01 |
| RG228 | USA | America | 1980 | American & European mixed | 0.14 | 0.09 | 0.77 |
| RG229 | Sweden | Europe | 1988 | European mixed | 0.46 | 0.48 | 0.06 |
| RG230 | Univ.Calif. | America | 1982 | American & European mixed | 0.16 | 0.37 | 0.48 |
| RG232 | France | Europe | 1900 | Heirloom & related | 1.00 | 0.00 | 0.00 |
| RG234 | Germany | Europe | 1954 | Heirloom & related | 0.59 | 0.31 | 0.11 |
| RG235 | Univ.Calif. | America | 1968 | American & European mixed | 0.12 | 0.26 | 0.62 |
| RG236 | NA | NA | 2001 | American & European mixed | 0.28 | 0.19 | 0.54 |
| RG237 | Italy | Europe | 2016 | American & European mixed | 0.00 | 0.41 | 0.59 |
| RG238 | Netherlands | Europe | 2005 | European mixed | 0.09 | 0.91 | 0.00 |
| RG240 | Univ.Calif. | America | 1982 | American & European mixed | 0.22 | 0.00 | 0.78 |
| RG241 | Belgium | Europe | 1942 | Heirloom & related | 0.89 | 0.11 | 0.00 |
| RG243 | Germany | Europe | 1922 | Heirloom & related | 0.91 | 0.00 | 0.09 |
| RG244 | France | Europe | 1896 | Heirloom & related | 0.77 | 0.21 | 0.02 |
| RG245 | Canada | America | 2004 | European mixed | 0.31 | 0.63 | 0.05 |
| RG246 | Canada | America | 2004 | European mixed | 0.20 | 0.75 | 0.05 |
| RG248 | Maryland | America | 1951 | European mixed | 0.15 | 0.85 | 0.00 |
| RG250 | Univ.Calif. | America | 1980 | European mixed | 0.31 | 0.37 | 0.32 |
| RG251 | California | America | 2012 | American & European mixed | 0.03 | 0.00 | 0.97 |
| RG253 | Italy | Europe | 2015 | European mixed | 0.02 | 0.63 | 0.35 |
| RG254 | Italy | Europe | 2000 | European mixed | 0.37 | 0.63 | 0.00 |
| RG256 | Univ.Calif. | America | 1964 | American & European mixed | 0.18 | 0.09 | 0.73 |
| RG258 | Japan | Asia | 1996 | European mixed | 0.28 | 0.44 | 0.28 |
| RG259 | Netherlands | Europe | 2013 | Heirloom & related | 0.40 | 0.28 | 0.32 |
| RG261 | Univ.Calif. | America | 1982 | American & European mixed | 0.06 | 0.01 | 0.93 |
| RG263 | Maryland | America | 1992 | European mixed | 0.18 | 0.76 | 0.06 |
| RG264 | Maryland | America | 1992 | European mixed | 0.15 | 0.68 | 0.18 |
| RG265 | Netherlands | Europe | 1983 | European mixed | 0.08 | 0.92 | 0.00 |
| RG266 | Univ.Calif. | America | 2001 | American & European mixed | 0.00 | 0.00 | 1.00 |
| RG269 | Netherlands | Europe | 1988 | European mixed | 0.07 | 0.81 | 0.12 |
| RG270 | Germany | Europe | 1963 | Heirloom & related | 0.51 | 0.44 | 0.05 |
| RG271 | France | Europe | 1922 | Heirloom & related | 0.83 | 0.08 | 0.10 |
| RG272 | France | Europe | 1929 | Heirloom & related | 0.84 | 0.16 | 0.00 |
| RG274 | Netherlands | Europe | 2007 | American & European mixed | 0.09 | 0.00 | 0.91 |
| RG275 | Italy | Europe | 2014 | American & European mixed | 0.00 | 0.20 | 0.80 |
| RG276 | Italy | Europe | 2000 | European mixed | 0.08 | 0.92 | 0.00 |
| RG277 | USA | America | 1860 | Heirloom & related | 0.89 | 0.05 | 0.05 |
| RG278 | UK | Europe | 1997 | European mixed | 0.46 | 0.49 | 0.05 |
| RG279 | Netherlands | Europe | 1998 | American & European mixed | 0.00 | 0.42 | 0.58 |

**Supplementary Table S2**. Pearson correlations between 2-year estimated BLUP values for the 12 traits.

| Trait 1 | Trait 2 | Pearson correlation | p-value |
|---|---|---|---|
| Skin Resistance | Bruiseness | 0.87 | 0 |
| Firmness | Bruiseness | 0.73 | 0 |
| Firmness | Skin Resistance | 0.72 | 0 |
| Color Homogeneity | Shape Homogeneity | 0.61 | 0 |
| Bruiseness | Glossyness | 0.60 | 0 |
| Mean Weight | Glossyness | 0.58 | 0 |
| Skin Resistance | Glossyness | 0.58 | 0 |
| Mean Weight | Firmness | 0.54 | 0 |
| Mean Weight | Skin Resistance | 0.51 | 1.51E-14 |
| Firmness | Glossyness | 0.50 | 1.91E-14 |
| Mean Weight | Bruiseness | 0.50 | 2.65E-13 |
| Glossyness | Color Homogeneity | 0.47 | 1.20E-12 |
| Titratable Acidity | Total Soluble Solids | 0.44 | 2.03E-11 |
| Glossyness | Shape Homogeneity | 0.44 | 5.23E-11 |
| Bruiseness | Color Homogeneity | 0.41 | 1.28E-09 |
| Skin Resistance | Color Homogeneity | 0.40 | 1.53E-09 |
| Skin Resistance | Shape Homogeneity | 0.40 | 1.91E-09 |
| Bruiseness | Shape Homogeneity | 0.39 | 1.30E-08 |
| Mean Weight | Color Homogeneity | 0.36 | 6.51E-08 |
| Firmness | Color Homogeneity | 0.35 | 1.22E-07 |
| Firmness | Shape Homogeneity | 0.34 | 3.88E-07 |
| Mean Weight | Shape Homogeneity | 0.29 | 2.29E-05 |
| Shape Homogeneity | Total Soluble Solids | 0.27 | 6.26E-05 |
| Color Homogeneity | Total Soluble Solids | 0.26 | 9.73E-05 |
| Firmness | External Color | 0.23 | 0.0008 |
| Skin Resistance | External Color | 0.20 | 0.00474 |
| Mean Weight | External Color | 0.17 | 0.01721 |
| Bruiseness | External Color | 0.16 | 0.02319 |
| Achene Insertion | Titratable Acidity | 0.11 | 0.12552 |
| Glossyness | External Color | 0.10 | 0.15265 |
| Color Homogeneity | Titratable Acidity | 0.10 | 0.1543 |
| Shape Homogeneity | Titratable Acidity | 0.09 | 0.17076 |
| Skin Resistance | Total Soluble Solids | 0.05 | 0.49268 |
| Firmness | Achene Insertion | 0.04 | 0.54422 |
| Mean Weight | Achene Insertion | 0.01 | 0.87498 |
| Glossyness | Total Soluble Solids | 0.01 | 0.91485 |
| Bruiseness | Total Soluble Solids | 0.00 | 0.95612 |
| Achene Insertion | External Color | -0.01 | 0.88308 |
| Skin Resistance | Titratable Acidity | -0.01 | 0.83638 |
| Achene Insertion | Total Soluble Solids | -0.02 | 0.74097 |
| Shape Homogeneity | External Color | -0.02 | 0.73203 |
| Titratable Acidity | External Color | -0.03 | 0.67376 |
| Color Homogeneity | External Color | -0.05 | 0.5183 |
| Bruiseness | Titratable Acidity | -0.05 | 0.48536 |
| Firmness | Total Soluble Solids | -0.07 | 0.29334 |
| Firmness | Titratable Acidity | -0.08 | 0.23537 |
| Achene Insertion | Color Homogeneity | -0.09 | 0.1992 |
| Achene Insertion | Shape Homogeneity | -0.09 | 0.18106 |
| Glossyness | Titratable Acidity | -0.11 | 0.13446 |
| Total Soluble Solids | External Color | -0.12 | 0.08283 |
| Bruiseness | Achene Insertion | -0.14 | 0.04316 |
| Skin Resistance | Achene Insertion | -0.14 | 0.0373 |
| Mean Weight | Titratable Acidity | -0.17 | 0.01697 |
| Glossyness | Achene Insertion | -0.21 | 0.00234 |
| Mean Weight | Total Soluble Solids | -0.23 | 0.00088 |

**Supplementary Table S3**. List of significant trait associations obtained for the GWAS on the 12 traits. Position Camarosa and Position Royal Royce: physical positions on Camarosa and Royal Royce reference genomes. MAF, minor allele frequency; PVE, Phenotypic variance explained (%).

| Trait | Year | SNP | Chromosome | Position Camarosa | Position Royal Royce | pval_Mahalanobis | threshold | π_group1 | π_group2 | π_group3 |
|---|---|---|---|---|---|---|---|---|---|---|
| Glossiness (GLO) | 2021 | AX-184177060 | 3D | 27845440 | 3815908 | 0.001 | ** | 0.44 | 0.32 | 0.15 |
| Glossiness (GLO) | 2020 | AX-184177060 | 3D | 27845440 | 3815908 | 0.001 | ** | 0.44 | 0.32 | 0.15 |
| Glossiness (GLO) | combinedvalues | AX-184177060 | 3D | 27845440 | 3815908 | 0.001 | ** | 0.44 | 0.32 | 0.15 |
| Skin resistance (SR) | 2021 | AX-184177060 | 3D | 27845440 | 3815908 | 0.001 | ** | 0.44 | 0.32 | 0.15 |
| Firmness (FIRM) | 2021 | AX-184477554 | 3D | 29275014 | 2454715 | 0.008 | ** | 0.40 | 0.39 | 0.29 |
| Shape homogeneity (UFS) | Combinedvalues | AX-184880676 | 2B | 25135835 | 25837267 | 0.008 | ** | 0.36 | 0.30 | 0.31 |
| Glossiness (GLO) | 2020 | AX-184352835 | 5A | 28239789 | 24966878 | 0.031 | * | 0.35 | 0.34 | 0.40 |
| Skin resistance (SR) | 2021 | AX-184130926 | 7A | 25616815 | 18667225 | 0.032 | * | 0.31 | 0.35 | 0.35 |
| TSS (Brix) | Combinedvalues | AX-184864732 | 6D | 12013420 | 9512125 | 0.041 | * | 0.36 | 0.43 | 0.22 |
| Titratable acidity (TA) | 2020 | AX-184457703 | 3D | 30090404 | -- | 0.066 | | 0.37 | 0.40 | 0.34 |
| Skin color (COL) | Combinedvalues | AX-184300087 | 7B | 1386945 | 23020879 | 0.084 | | 0.34 | 0.33 | 0.30 |
| Achene position (ACH) | Combinedvalues | AX-184381485 | 2C | 19190988 | 7710274 | 0.088 | | 0.39 | 0.36 | 0.37 |
| TSS (Brix) | Combinedvalues | AX-184505625 | 3B | 4434681 | 4139234 | 0.138 | | 0.39 | 0.44 | 0.39 |
| Skin resistance (SR) | 2021 | AX-123359788 | 7A | 28581204 | -- | 0.139 | | 0.31 | 0.34 | 0.34 |
| TSS (Brix) | Combinedvalues | AX-184920058 | 7B | 4515979 | 19984301 | 0.179 | | 0.41 | 0.43 | 0.30 |
| TSS (Brix) | 2020 | AX-184920058 | 7B | 4515979 | 19984301 | 0.179 | | 0.41 | 0.43 | 0.30 |
| TSS (Brix) | Combinedvalues | AX-184560339 | 7C | 19112988 | 13209597 | 0.234 | | 0.42 | 0.34 | 0.35 |
| Titratable acidity (TA) | Combinedvalues | AX-166514185 | 5C | 6983698 | 8203430 | 0.238 | | 0.34 | 0.37 | 0.34 |
| Glossiness (GLO) | 2021 | AX-184494194 | 1C | 11470545 | 10966861 | 0.253 | | 0.43 | 0.42 | 0.18 |
| Titratable acidity (TA) | Combinedvalues | AX-184595531 | 6A | 25621066 | 9172070 | 0.301 | | 0.36 | 0.37 | 0.41 |
| Achene position (ACH) | Combinedvalues | AX-184310362 | 6D | 939082 | 32008538 | 0.31 | | 0.38 | 0.40 | 0.30 |
| TSS (Brix) | 2020 | AX-184355355 | 6B | 27647943 | 10765128 | 0.313 | | 0.37 | 0.34 | 0.34 |
| Skin color (COL) | Combinedvalues | AX-166514401 | 5C | 11987143 | -- | 0.351 | | 0.35 | 0.37 | 0.39 |
| TSS (Brix) | 2020 | AX-184131652 | 1B | 4174690 | 1302571 | 0.358 | | 0.25 | 0.36 | 0.39 |
| Bruiseness (BRU) | combinedvalues | AX-184399480 | 6B | 5731875 | 30532752 | 0.364 | | 0.41 | 0.33 | 0.43 |
| Glossiness (GLO) | combinedvalues | AX-184408294 | 5A | 28658239 | 25341723 | 0.39 | | 0.35 | 0.36 | 0.44 |
| Achene position (ACH) | Combinedvalues | AX-184747287 | 6C | 32305891 | 30352992 | 0.393 | | 0.42 | 0.45 | 0.38 |
| TSS (Brix) | Combinedvalues | AX-184970304 | 5A | 12400711 | 10544768 | 0.396 | | 0.32 | 0.41 | 0.36 |
| TSS (Brix) | 2020 | AX-184970304 | 5A | 12400711 | 10544768 | 0.396 | | 0.32 | 0.41 | 0.36 |
| Shape homogeneity (UFS) | Combinedvalues | AX-184043005 | 2B | 11149178 | 11940989 | 0.413 | | 0.38 | 0.42 | 0.31 |
| Shape homogeneity (UFS) | Combinedvalues | AX-184466777 | 2D | 12871717 | 11774978 | 0.413 | | 0.35 | 0.42 | 0.41 |
| Firmness (FIRM) | 2021 | AX-184039356 | 6A | 7277130 | 27533782 | 0.415 | | 0.34 | 0.38 | 0.36 |
| Glossiness (GLO) | combinedvalues | AX-184951955 | 4C | 727143 | 26073661 | 0.436 | | 0.38 | 0.35 | 0.37 |
| TSS (Brix) | 2020 | AX-184330352 | 5A | 22141939 | 18780692 | 0.472 | | 0.31 | 0.45 | 0.38 |
| Firmness (FIRM) | 2021 | AX-123522331 | 2C | 22908001 | 3685634 | 0.475 | | 0.34 | 0.36 | 0.39 |
| Titratable acidity (TA) | 2020 | AX-184091372 | 1A | 13540517 | 13469152 | 0.489 | | 0.38 | 0.39 | 0.39 |
| TSS (Brix) | 2020 | AX-184221848 | 6C | 33293635 | 29374007 | 0.499 | | 0.43 | 0.46 | 0.33 |
| TSS (Brix) | 2020 | AX-184507945 | 5A | 21971093 | 18609839 | 0.5 | | 0.33 | 0.45 | 0.37 |
| TSS (Brix) | 2020 | AX-184219801 | 5A | 22266307 | 18905072 | 0.5 | | 0.31 | 0.45 | 0.38 |
| Glossiness (GLO) | combinedvalues | AX-184599570 | 3D | 26901693 | 4775280 | 0.509 | | 0.39 | 0.39 | 0.21 |
| Skin color (COL) | Combinedvalues | AX-184090167 | 7C | 19438028 | 13528856 | 0.535 | | 0.35 | 0.35 | 0.34 |
| TSS (Brix) | 2020 | AX-184079508 | 7C | 26901718 | 19354380 | 0.545 | | 0.32 | 0.34 | 0.38 |
| TSS (Brix) | Combinedvalues | AX-184179821 | 5A | 22384778 | 18999088 | 0.553 | | 0.31 | 0.45 | 0.38 |
| Skin color (COL) | 2020 | AX-184965421 | 5D | 14022053 | 13542145 | 0.555 | | 0.35 | 0.43 | 0.41 |
| Skin color (COL) | Combinedvalues | AX-184965421 | 5D | 14022053 | 13542145 | 0.555 | | 0.35 | 0.43 | 0.41 |
| Bruiseness (BRU) | combinedvalues | AX-184713611 | 7C | 20943204 | 15149370 | 0.56 | | 0.41 | 0.41 | 0.39 |
| Titratable acidity (TA) | Combinedvalues | AX-184462338 | 6C | 27458040 | -- | 0.575 | | 0.35 | 0.41 | 0.39 |
| Bruiseness (BRU) | combinedvalues | AX-184940044 | 1A | 3531536 | -- | 0.623 | | 0.28 | 0.39 | 0.42 |
| Fruit weight (FW) | combinedvalues | AX-184413183 | 1B | 19119571 | 15971709 | 0.639 | | 0.38 | 0.41 | 0.35 |
| Firmness (FIRM) | 2021 | AX-184521799 | 2A | 17814521 | 5400298 | 0.641 | | 0.36 | 0.38 | 0.41 |
| B/Aratio | 2021 | AX-184452909 | 6D | 1646690 | 32715821 | 0.685 | | 0.44 | 0.41 | 0.32 |
| TSS (Brix) | 2020 | AX-184477629 | 3B | 1541393 | 1842542 | 0.757 | | 0.37 | 0.43 | 0.38 |
| Shape homogeneity (UFS) | Combinedvalues | AX-184477629 | 3B | 1541393 | 1842542 | 0.757 | | 0.37 | 0.43 | 0.38 |
| TSS (Brix) | 2020 | AX-184418966 | 5A | 24950308 | 21580601 | 0.775 | | 0.29 | 0.39 | 0.41 |
| B/Aratio | Combinedvalues | AX-184399755 | 6B | 31578303 | 9217798 | 0.793 | | 0.39 | 0.27 | 0.20 |
| TSS (Brix) | 2021 | AX-184399755 | 6B | 31578303 | 9217798 | 0.793 | | 0.39 | 0.27 | 0.20 |
| Skin color (COL) | Combinedvalues | AX-184965919 | 6A | 14476176 | 21070121 | 0.799 | | 0.29 | 0.36 | 0.38 |
| Skin resistance (SR) | 2021 | AX-184230747 | 5B | 15853776 | 12173853 | 0.804 | | 0.35 | 0.39 | 0.41 |
| Bruiseness (BRU) | combinedvalues | AX-184408176 | 5A | 18814640 | 16414924 | 0.838 | | 0.30 | 0.34 | 0.31 |
| Fruit weight (FW) | combinedvalues | AX-184592155 | 2D | 15565564 | 8801569 | 0.876 | | 0.40 | 0.44 | 0.41 |
| Shape homogeneity (UFS) | 2021 | AX-184458801 | 3A | 21213134 | 9650845 | 0.88 | | 0.30 | 0.36 | 0.43 |
| Skin resistance (SR) | 2021 | AX-184127736 | 4A | 20012930 | 16433479 | 0.916 | | 0.36 | 0.34 | 0.34 |
| Fruit weight (FW) | combinedvalues | AX-184241601 | 5B | 17045086 | 10918733 | 0.925 | | 0.32 | 0.30 | 0.34 |
| B/Aratio | 2021 | AX-184052133 | 3B | 4097709 | 3828069 | 0.937 | | 0.34 | 0.39 | 0.35 |
| Bruiseness (BRU) | combinedvalues | AX-184203769 | 1B | 9439391 | 7276991 | 0.954 | | 0.32 | 0.37 | 0.44 |
| TSS (Brix) | 2021 | AX-184047575 | 7C | 20267768 | -- | 0.957 | | 0.41 | 0.40 | 0.37 |
| B/Aratio | 2021 | AX-184282016 | 3B | 4095262 | 3825622 | 0.968 | | 0.34 | 0.39 | 0.35 |
| TSS (Brix) | 2020 | AX-184718481 | 2D | 13298291 | -- | 0.982 | | 0.40 | 0.46 | 0.41 |
| Shape homogeneity (UFS) | 2021 | AX-184554177 | 1A | 10002081 | 10295193 | 0.994 | | 0.30 | 0.34 | 0.35 |
| Shape homogeneity (UFS) | 2021 | AX-184611387 | 1A | 9927298 | 10221629 | 0.998 | | 0.31 | 0.42 | 0.42 |
| Shape homogeneity (UFS) | 2021 | AX-89904139 | 1A | 9957207 | -- | 0.998 | | 0.30 | 0.34 | 0.35 |

**Supplementary Table S4**. Genome scan outputs for the 71 trait associations. Position Camarosa and Position Royal Royce: physical positions on Camarosa and Royal Royce reference genomes. pval_Mahalanobis, p-values of the genome scan based on Mahalanobis distance. threshold, p-value thresholds at 0.01 (**) and 0.05 (*). π_group1, π_group2, π_group3, π values of the genome scans performed on respectively genetic groups 1, 2 and 3 within 400kb windows.

| Trait | Year | SNP | Chromosome | Position Camarosa | Position Royal Royce | pval_Mahalanobis | threshold | π_group1 | π_group2 | π_group3 |
|---|---|---|---|---|---|---|---|---|---|---|
| Glossiness (GLO) | 2021 | AX-184177060 | 3D | 27845440 | 3815908 | 0.001 | ** | 0.44 | 0.32 | 0.15 |
| Glossiness (GLO) | 2020 | AX-184177060 | 3D | 27845440 | 3815908 | 0.001 | ** | 0.44 | 0.32 | 0.15 |
| Glossiness (GLO) | combinedvalues | AX-184177060 | 3D | 27845440 | 3815908 | 0.001 | ** | 0.44 | 0.32 | 0.15 |
| Skin resistance (SR) | 2021 | AX-184177060 | 3D | 27845440 | 3815908 | 0.001 | ** | 0.44 | 0.32 | 0.15 |
| Firmness (FIRM) | 2021 | AX-184477554 | 3D | 29275014 | 2454715 | 0.008 | ** | 0.40 | 0.39 | 0.29 |
| Shape homogeneity (UFS) | Combinedvalues | AX-184880676 | 2B | 25135835 | 25837267 | 0.008 | ** | 0.36 | 0.30 | 0.31 |
| Glossiness (GLO) | 2020 | AX-184352835 | 5A | 28239789 | 24966878 | 0.031 | * | 0.35 | 0.34 | 0.40 |
| Skin resistance (SR) | 2021 | AX-184130926 | 7A | 25616815 | 18667225 | 0.032 | * | 0.31 | 0.35 | 0.35 |
| TSS (Brix) | Combinedvalues | AX-184864732 | 6D | 12013420 | 9512125 | 0.041 | * | 0.36 | 0.43 | 0.22 |
| Titratable acidity (TA) | 2020 | AX-184457703 | 3D | 30090404 | -- | 0.066 | | 0.37 | 0.40 | 0.34 |
| Skin color (COL) | Combinedvalues | AX-184300087 | 7B | 1386945 | 23020879 | 0.084 | | 0.34 | 0.33 | 0.30 |
| Achene position (ACH) | Combinedvalues | AX-184381485 | 2C | 19190988 | 7710274 | 0.088 | | 0.39 | 0.36 | 0.37 |
| TSS (Brix) | Combinedvalues | AX-184505625 | 3B | 4434681 | 4139234 | 0.138 | | 0.39 | 0.44 | 0.39 |
| Skin resistance (SR) | 2021 | AX-123359788 | 7A | 28581204 | -- | 0.139 | | 0.31 | 0.34 | 0.34 |
| TSS (Brix) | Combinedvalues | AX-184920058 | 7B | 4515979 | 19984301 | 0.179 | | 0.41 | 0.43 | 0.30 |
| TSS (Brix) | 2020 | AX-184920058 | 7B | 4515979 | 19984301 | 0.179 | | 0.41 | 0.43 | 0.30 |
| TSS (Brix) | Combinedvalues | AX-184560339 | 7C | 19112988 | 13209597 | 0.234 | | 0.42 | 0.34 | 0.35 |
| Titratable acidity (TA) | Combinedvalues | AX-166514185 | 5C | 6983698 | 8203430 | 0.238 | | 0.34 | 0.37 | 0.34 |
| Glossiness (GLO) | 2021 | AX-184494194 | 1C | 11470545 | 10966861 | 0.253 | | 0.43 | 0.42 | 0.18 |
| Titratable acidity (TA) | Combinedvalues | AX-184595531 | 6A | 25621066 | 9172070 | 0.301 | | 0.36 | 0.37 | 0.41 |
| Achene position (ACH) | Combinedvalues | AX-184310362 | 6D | 939082 | 32008538 | 0.31 | | 0.38 | 0.40 | 0.30 |
| TSS (Brix) | 2020 | AX-184355355 | 6B | 27647943 | 10765128 | 0.313 | | 0.37 | 0.34 | 0.34 |
| Skin color (COL) | Combinedvalues | AX-166514401 | 5C | 11987143 | -- | 0.351 | | 0.35 | 0.37 | 0.39 |
| TSS (Brix) | 2020 | AX-184131652 | 1B | 4174690 | 1302571 | 0.358 | | 0.25 | 0.36 | 0.39 |
| Bruiseness (BRU) | combinedvalues | AX-184399480 | 6B | 5731875 | 30532752 | 0.364 | | 0.41 | 0.33 | 0.43 |
| Glossiness (GLO) | combinedvalues | AX-184408294 | 5A | 28658239 | 25341723 | 0.39 | | 0.35 | 0.36 | 0.44 |
| Achene position (ACH) | Combinedvalues | AX-184747287 | 6C | 32305891 | 30352992 | 0.393 | | 0.42 | 0.45 | 0.38 |
| TSS (Brix) | Combinedvalues | AX-184970304 | 5A | 12400711 | 10544768 | 0.396 | | 0.32 | 0.41 | 0.36 |
| TSS (Brix) | 2020 | AX-184970304 | 5A | 12400711 | 10544768 | 0.396 | | 0.32 | 0.41 | 0.36 |
| Shape homogeneity (UFS) | Combinedvalues | AX-184043005 | 2B | 11149178 | 11940989 | 0.413 | | 0.38 | 0.42 | 0.31 |
| Shape homogeneity (UFS) | Combinedvalues | AX-184466777 | 2D | 12871717 | 11774978 | 0.413 | | 0.35 | 0.42 | 0.41 |
| Firmness (FIRM) | 2021 | AX-184039356 | 6A | 7277130 | 27533782 | 0.415 | | 0.34 | 0.38 | 0.36 |
| Glossiness (GLO) | combinedvalues | AX-184951955 | 4C | 727143 | 26073661 | 0.436 | | 0.38 | 0.35 | 0.37 |
| TSS (Brix) | 2020 | AX-184330352 | 5A | 22141939 | 18780692 | 0.472 | | 0.31 | 0.45 | 0.38 |
| Firmness (FIRM) | 2021 | AX-123522331 | 2C | 22908001 | 3685634 | 0.475 | | 0.34 | 0.36 | 0.39 |
| Titratable acidity (TA) | 2020 | AX-184091372 | 1A | 13540517 | 13469152 | 0.489 | | 0.38 | 0.39 | 0.39 |
| TSS (Brix) | 2020 | AX-184221848 | 6C | 33293635 | 29374007 | 0.499 | | 0.43 | 0.46 | 0.33 |
| TSS (Brix) | 2020 | AX-184507945 | 5A | 21971093 | 18609839 | 0.5 | | 0.33 | 0.45 | 0.37 |
| TSS (Brix) | 2020 | AX-184219801 | 5A | 22266307 | 18905072 | 0.5 | | 0.31 | 0.45 | 0.38 |
| Glossiness (GLO) | combinedvalues | AX-184599570 | 3D | 26901693 | 4775280 | 0.509 | | 0.39 | 0.39 | 0.21 |
| Skin color (COL) | Combinedvalues | AX-184090167 | 7C | 19438028 | 13528856 | 0.535 | | 0.35 | 0.35 | 0.34 |
| TSS (Brix) | 2020 | AX-184079508 | 7C | 26901718 | 19354380 | 0.545 | | 0.32 | 0.34 | 0.38 |
| TSS (Brix) | Combinedvalues | AX-184179821 | 5A | 22384778 | 18999088 | 0.553 | | 0.31 | 0.45 | 0.38 |
| Skin color (COL) | 2020 | AX-184965421 | 5D | 14022053 | 13542145 | 0.555 | | 0.35 | 0.43 | 0.41 |
| Skin color (COL) | Combinedvalues | AX-184965421 | 5D | 14022053 | 13542145 | 0.555 | | 0.35 | 0.43 | 0.41 |
| Bruiseness (BRU) | combinedvalues | AX-184713611 | 7C | 20943204 | 15149370 | 0.56 | | 0.41 | 0.41 | 0.39 |
| Titratable acidity (TA) | Combinedvalues | AX-184462338 | 6C | 27458040 | -- | 0.575 | | 0.35 | 0.41 | 0.39 |
| Bruiseness (BRU) | combinedvalues | AX-184940044 | 1A | 3531536 | -- | 0.623 | | 0.28 | 0.39 | 0.42 |
| Fruit weight (FW) | combinedvalues | AX-184413183 | 1B | 19119571 | 15971709 | 0.639 | | 0.38 | 0.41 | 0.35 |
| Firmness (FIRM) | 2021 | AX-184521799 | 2A | 17814521 | 5400298 | 0.641 | | 0.36 | 0.38 | 0.41 |
| B/Aratio | 2021 | AX-184452909 | 6D | 1646690 | 32715821 | 0.685 | | 0.44 | 0.41 | 0.32 |
| TSS (Brix) | 2020 | AX-184477629 | 3B | 1541393 | 1842542 | 0.757 | | 0.37 | 0.43 | 0.38 |
| Shape homogeneity (UFS) | Combinedvalues | AX-184477629 | 3B | 1541393 | 1842542 | 0.757 | | 0.37 | 0.43 | 0.38 |
| TSS (Brix) | 2020 | AX-184418966 | 5A | 24950308 | 21580601 | 0.775 | | 0.29 | 0.39 | 0.41 |
| B/Aratio | Combinedvalues | AX-184399755 | 6B | 31578303 | 9217798 | 0.793 | | 0.39 | 0.27 | 0.20 |
| TSS (Brix) | 2021 | AX-184399755 | 6B | 31578303 | 9217798 | 0.793 | | 0.39 | 0.27 | 0.20 |
| Skin color (COL) | Combinedvalues | AX-184965919 | 6A | 14476176 | 21070121 | 0.799 | | 0.29 | 0.36 | 0.38 |
| Skin resistance (SR) | 2021 | AX-184230747 | 5B | 15853776 | 12173853 | 0.804 | | 0.35 | 0.39 | 0.41 |
| Bruiseness (BRU) | combinedvalues | AX-184408176 | 5A | 18814640 | 16414924 | 0.838 | | 0.30 | 0.34 | 0.31 |
| Fruit weight (FW) | combinedvalues | AX-184592155 | 2D | 15565564 | 8801569 | 0.876 | | 0.40 | 0.44 | 0.41 |
| Shape homogeneity (UFS) | 2021 | AX-184458801 | 3A | 21213134 | 9650845 | 0.88 | | 0.30 | 0.36 | 0.43 |
| Skin resistance (SR) | 2021 | AX-184127736 | 4A | 20012930 | 16433479 | 0.916 | | 0.36 | 0.34 | 0.34 |
| Fruit weight (FW) | combinedvalues | AX-184241601 | 5B | 17045086 | 10918733 | 0.925 | | 0.32 | 0.30 | 0.34 |
| B/Aratio | 2021 | AX-184052133 | 3B | 4097709 | 3828069 | 0.937 | | 0.34 | 0.39 | 0.35 |
| Bruiseness (BRU) | combinedvalues | AX-184203769 | 1B | 9439391 | 7276991 | 0.954 | | 0.32 | 0.37 | 0.44 |
| TSS (Brix) | 2021 | AX-184047575 | 7C | 20267768 | -- | 0.957 | | 0.41 | 0.40 | 0.37 |
| B/Aratio | 2021 | AX-184282016 | 3B | 4095262 | 3825622 | 0.968 | | 0.34 | 0.39 | 0.35 |
| TSS (Brix) | 2020 | AX-184718481 | 2D | 13298291 | -- | 0.982 | | 0.40 | 0.46 | 0.41 |
| Shape homogeneity (UFS) | 2021 | AX-184554177 | 1A | 10002081 | 10295193 | 0.994 | | 0.30 | 0.34 | 0.35 |
| Shape homogeneity (UFS) | 2021 | AX-184611387 | 1A | 9927298 | 10221629 | 0.998 | | 0.31 | 0.42 | 0.42 |
| Shape homogeneity (UFS) | 2021 | AX-89904139 | 1A | 9957207 | -- | 0.998 | | 0.30 | 0.34 | 0.35 |

Supplementary Table S5. Functions and/or possible roles of the 64 candidate genes underlying the trait associations.

| Trait | Chromosome | Protein encoded by the Candidate Gene (CG) | *CG Abbreviation* | Function and/or possible role |
|---|---|---|---|---|
| Fruit weight | 1B | cyclin-dependent kinase E-1 | *CDKE* | cell division and regulation of developmental growth |
| | 2D | small auxin upregulated RNA 14 | *SAUR14* | cell differentiation and regulation of developmental growth |
| | | small auxin upregulated RNA 1 | *SAUR1* | cell differentiation and regulation of developmental growth |
| | | small auxin upregulated RNA 20 | *SAUR20* | cell differentiation and regulation of developmental growth |
| | | small auxin upregulated RNA 51 | *SAUR51* | cell differentiation and regulation of developmental growth |
| | | small auxin upregulated RNA 49 | *SAUR49* | cell differentiation and regulation of developmental growth |
| | 5B | cullin | *CUL* | a component of SCF ubiquitin ligase complexes involved in mediating responses to auxin |
| | | cullin | *CUL* | a component of SCF ubiquitin ligase complexes involved in mediating responses to auxin |
| Skin color | 5C | anthocyanidin 3-O-glucosyltransferase | *FaGT2* | glycosyltransferase that transfers UDP-glucose to anthocyanidins thus generating stable 3-O-glucosides |
| | 5D | flavonoid 3-O-glycosyltransferase | *GT* | glycosyltransferase that transfers UDP-glucose to anthocyanidins thus generating stable 3-*O*-glucosides |
| | 6A | caffeoylshikimate esterase | *CSE* | key enzyme in the lignin biosynthetic pathway |
| | 7B | anthocyanidin 3-O-glucosyltransferase | *FaGT1* | glycosyltransferase that transfers UDP-glucose to anthocyanidins thus generating stable 3-*O*-glucosides |
| | 7C | TT12-like MATE transporter | *TT12* | transport of anthocyanidins to the vacuole |
| Firmness | 3D | AGP galactosyltransferase | *GALT* | hydroxyproline-O-galactosyltransferase specific for cell wall arabinogalactan-protein biosynthesis |
| | | * xyloglucan endotransglucosylase/hydrolase | *XTH* | catalyzes the cleavage of xyloglucans, thus functions in the loosening and rearrangement of the cell wall |
| | | * cellulose synhase | *CES* | involved in cellulose synthesis and cell wall formation |
| | 6A | cellulase 1 | *CEL* | endo-1,4-beta-glucanase activity, involved in cell wall modifications and cell elongation |
| | | polygalacturonase | *PG* | polygalacturonase activity that depolymerizes cell wall pectins; involved in strawberry fruit softening |
| | | polygalacturonase | *PG* | polygalacturonase activity that depolymerizes cell wall pectins; involved in strawberry fruit softening |
| Titratable acid | 1A | pyruvate kinase | *PK* | involved in glycolysis and in the control of citric acid during strawberry fruit ripening |
| | 6A | V-type proton ATPase subunit G | *VMA-G* | vacuolar ATPase establishes the electrochemical gradient for proton across the tonoplast |
| | 6C | V-type proton ATPase subunit C | *VMA-C* | vacuolar ATPase establishes the electrochemical gradient for proton across the tonoplast |
| TSS | 1B | sucrose-phosphate synthase | *SPS* | involved in sucrose synthesis |
| | | sucrose-phosphate synthase | *SPS* | involved in sucrose synthesis |
| | 2D | fructose-1,6-bisphosphatase, cytosolic | *FBP* | formation of fructose-6-phosphate for sucrose biosynthesis, possible role in fructose-mediated signaling |
| | 3B | * starch synthase | *SS* | involved in starch granule initiation and formation |
| | 5A | fructose-1,6-bisphosphatase, cytosolic | *FBP* | formation of fructose-6-phosphate for sucrose biosynthesis, possible role in fructose-mediated signaling |
| | 6B | fructose-bisphosphate aldolase | *FBA* | a key metabolic enzyme in glycolysis and gluconeogenesis in plants |
| | | isocitrate dehydrogenase [NAD] | *IDH* | produces 2-oxoglutarate; key regulatory step of the TCA cycle |
| | 6C | aconitase | *ACO* | can catalyze the conversion of citrate to isocitrate, may participate in primary metabolism including the TCA cycle |
| | 6D | isocitrate dehydrogenase [NAD] | *IDH* | produces 2-oxoglutarate; key regulatory step of the TCA cycle |
| | 7B | hexose carrier protein 6 | *HEX* | transport of hexoses |
| | 7C | alkaline/neutral invertase | *INV* | involved in sucrose breakdown to produce glucose and fructose |
| | | hexose carrier protein 6 | *HEX* | transport of hexoses |
| B/A ratio | 6B | isocitrate dehydrogenase [NAD] | *IDH* | produces 2-oxoglutarate; key regulatory step of the TCA cycle |
| Glossiness | 1C | cinnamyl-alcohol dehydrogenase | *CAD* | lignin biosynthesis |
| | | cinnamyl-alcohol dehydrogenase | *CAD* | lignin biosynthesis |
| | 3D | MYB-SHAQKYF | *MYS* | TF involved in wax biosynthesis; acts upstream of the DEWAX-SPL9 module with MYS2, thus regulating CER1 |
| | | trichome birefringence-like 38 | *TBL* | involved in epiderm differentiation, CW modifications, pectin esterification |
| | | * hydroxycinnamoyl-CoA shikimate/quinate hydroxycinnamoyltransferase | *HCT* | plays a critical function in the phenylpropanoid pathway in plants; is required for the formation of an intact cuticle |
| | | * GDSL esterase/lipase | *GELP* | large gene family with several members involved in the synthesis and depolymerisation of cutin and suberin |
| | | * GDSL esterase/lipase | *GELP* | large gene family with several members involved in the synthesis and depolymerisation of cutin and suberin |
| | | * GDSL esterase/lipase | *GELP* | large gene family with several members involved in the synthesis and depolymerisation of cutin and suberin |
| | 4C | non specific Lipid Transport Protein | *nsLTP* | possible transport of cuticle lipid precursors |
| | | glycerol-3-phosphate acyltransferase | *GPAT6* | synthesis of cutin precursors |
| | 5A | non specific Lipid Transport Protein | *nsLTP* | possible transport of cuticle lipid precursors |
| | | glycerol-3-phosphate acyltransferase | *GPAT3* | synthesis of cutin precursors |
| Skin resistance | 3D | MYB-SHAQKYF 1 | *MYS* | TF involved in wax biosynthesis; acts upstream of the DEWAX-SPL9 module with MYS2, thus regulating CER1 |
| | | trichome birefringence-like 38 | *TBL* | involved in epiderm differentiation, CW modifications, pectin esterification |
| | | * hydroxycinnamoyl-CoA shikimate/quinate hydroxycinnamoyltransferase | *HCT* | plays a critical function in the phenylpropanoid pathway in plants; required for the formation of an intact cuticle |
| | | * GDSL esterase/lipase | *GELP* | large gene family with several members involved in the assembly and depolymerisation of cutin and suberin |
| | | * GDSL esterase/lipase | *GELP* | large gene family with several members involved in the assembly and depolymerisation of cutin and suberin |
| | | * GDSL esterase/lipase | *GELP* | large gene family with several members involved in the assembly and depolymerisation of cutin and suberin |
| | 4A | COBRA-like | *COBL* | GPI anchored protein; key regulator of cellulose crystallinity and cell expansion |
| | | COBRA-like | *COBL* | GPI anchored protein; key regulator of cellulose crystallinity and cell expansion |
| | | beta-D-xylosidase | *BXL* | required for arabinan modification in CW pectins |
| | | trichome birefringence-like 43 | *TBL* | involved in secondary wall cellulose deposition, presumably through the esterification state of pectic polymers |
| | 5B | pectin methylesterase inhibitor | *PMEI* | inhibit pectin methylesterase activity thus leading to a higher degree of methylesterification of pectin |
| | 7A | polyglacturonase | *PG* | polygalacturonase activity that depolymerizes cell wall pectins |
| | | expansin B2 | *EXP* | mediates cell wall loosening and therefore modulates cell wall strength and structural integrity |
| | | expansin B2 | *EXP* | mediates cell wall loosening and therefore modulates cell wall strength and structural integrity |
| | | decrease was biosynthesis 2 | *DEWAX* | DEWAX2 Transcription Factor Negatively Regulates Cuticular Wax Biosynthesis in Arabidopsis Leaves |
| | | decrease was biosynthesis 2 | *DEWAX* | DEWAX2 Transcription Factor Negatively Regulates Cuticular Wax Biosynthesis in Arabidopsis Leaves |
| | 7A | wax ester synthase | *WSD* | involved in wax metabolism |
| Bruiseness | 1A | epidermal patterning factor | *EPFL* | plant specific secretory peptides with roles in the control of patterning in the plant epidermis |
| | 1B | GDSL lipase | *GELP* | large gene family with several members involved in the assembly and depolymerisation of cutin and suberin |
| | 5A | pectate lyase | *PL* | cleaves pectin homogalacturonan - involved in strawberry fruit softening |
| | 6B | fasciclin-like arabinogalactan protein | *FLA* | involved in cell wall formation and architecture |
| | | fasciclin-like arabinogalactan protein | *FLA* | involved in cell wall formation and architecture |
| | | xyloglucan endotransglucosylase/hydrolase | *XTH* | catalyzes the cleavage of xyloglucans, thus functions in the loosening and rearrangement of the cell wall |
| | 7C | 3-ketoacyl-CoA synthase 1-like | *KCS* | involved in cuticular wax synthesis; possibly involved in fruit water-loss after harvest |
